# Click-chemistry enabled directed evolution of glycosynthases for bespoke glycans synthesis

**DOI:** 10.1101/2020.03.23.001982

**Authors:** Ayushi Agrawal, Chandra Kanth Bandi, Tucker Burgin, Youngwoo Woo, Heather B. Mayes, Shishir P. S. Chundawat

## Abstract

Engineering of carbohydrate-active enzymes like glycosynthases for chemoenzymatic synthesis of bespoke oligosaccharides has been limited by the lack of suitable directed evolution based protein engineering methods. Currently there are no ultrahigh-throughput screening methods available for rapid and highly sensitive single cell-based screening of evolved glycosynthase enzymes employing azido sugars as substrates. Here, we report a fluorescence-based approach employing click-chemistry for the selective detection of glycosyl azides (versus free inorganic azides) that facilitated ultrahigh-throughput *in-vivo* single cell-based assay of glycosynthase activity. This discovery has led to the development of a directed evolution methodology for screening and sorting glycosynthase mutants for synthesis of desired fucosylated oligosaccharides. Our screening technique facilitated rapid fluorescence activated cell sorting of a large library of glycosynthase variants (>10^6^ mutants) expressed in *E. coli* to identify several novel mutants with increased activity for β-fucosyl-azide activated donor sugars towards desired acceptor sugars, demonstrating the broader applicability of this methodology.

## Introduction

Glycans are ubiquitous carbohydrate-based biomolecules found in nearly all living systems and are also the most abundant class of organic molecules on the planet, but we still have a limited understanding of their complex structural and functional roles in biology. One of the major stumbling blocks to advance the field of glycobiology identified has been the extreme difficulty in achieving the desired regio- and stereo-selectivity during synthesis of complex carbohydrate-based molecules using standard chemical or biochemical routes.^1^ Traditional methods for *de novo* synthesis of complex glycan polymers from monosaccharide building blocks requires complex reaction schemes with multiple protection-deprotection steps that often results in atom-inefficient product yields.^2,3^ This has limited the widespread adoption of chemical synthesis based approaches alone for commercial-scale applications. In contrast, enzymes can give high biosynthesis yields of glycans with the desired stereo- and regioselectivity.^4,5^ Glycosyltransferases (GTs) are primarily responsible for the natural biosynthesis of glycans facilitated by the transfer of nucleotide-glycosyl donors to targeted glycone or aglycone acceptors.^6^ However, GTs are typically membrane bound proteins that express poorly in standard protein expression systems like *Escherichia coli*, have limited solubility/stability, have high specificity that limits broader synthetic applications, and require recycling of expensive nucleotide donor sugar reagents that further limits scaling up *in vitro* biosynthetic routes.^7^ *In vivo* synthesis can partly address some of these challenges by either transferring or modifying the glycosylation biosynthetic pathways from desired eukaryotic or prokaryotic systems, like *Campylobacter jejuni*, into genetically tractable and industrially relevant expression systems like *E. coli* or *Pichia pastoris*.^8–10^ However, *in vivo* glycosylation is highly species-specific and produces a complex milieu of products due to the compounding presence of native carbohydrate-active enzymes (CAZymes) and/or other competing metabolic reactions that also utilize nucleotide sugars as substrates. It is thus challenging to synthesize structurally complex glycans using such chemical or biochemical based synthesis approaches alone. In recent years, there has been therefore an increased emphasis on the application of novel chemical-biology based approaches to accelerate the discovery of hybrid chemoenzymatic routes for glycans synthesis.^2^

Recent strides towards using glycosyl hydrolases (GHs) for *in vitro* chemoenzymatic synthesis has allowed production of bespoke complex glycans at high yields.^11^ GHs are nature’s antipode of GTs that catalyze the hydrolysis of glycosidic linkages, but can also synthesize glycans via the transglycosylation mechanism if the nucleophilic water is replaced by a suitable glycone or aglycone acceptor group instead. Unlike GTs, there is a larger selection of well-characterized GHs currently available that are highly soluble/stable, express readily in *E. coli*, and have the required structural plasticity for broadening substrate specificity using suitable protein engineering approaches, which overall makes GHs attractive biocatalysts for glycan synthesis. GHs are grouped into various families by amino-acid sequence similarity and substrate specificity,^12^ now numbering 166+ families as curated on the Carbohydrate-Active enZyme (CAZy) database.^13,14^ Unfortunately, transglycosylation reactions also suffer from low yields since the transglycosylation product is also the substrate for GHs. A promising solution to this issue was first pioneered by Withers and several others in 1998 through the application of a new class of mutant CAZymes called glycosynthases (GSs).^15–17^ These engineered enzymes are often single-point active-site mutants of native GHs to remove their ability to hydrolyze glycosidic bonds, but retain their ability to catalyze glycosidic bond formation between activated glycosyl donors like glycosyl fluorides (or glycosyl azides) and suitable acceptor groups to synthesize bespoke glycans. Mutation of the catalytic nucleophile residue (e.g., glutamate to alanine) often facilitates the transglycosylation reaction in the presence of activated sugar donors that mimic the glycosyl-enzyme intermediate (GEI) configuration, but while preventing hydrolysis of the transglycosylation product to give near-theoretical transglycosylation yields unlike wild-type GHs.^18^ However, there is not yet a truly rational approach, based on a comprehensive mechanistic understanding, that facilitates *in silico* design and engineering of GSs to produce bespoke oligosaccharides. Furthermore, there has been limited application of advanced protein engineering approaches to engineer GSs, which could explain why only a few successful GSs have been reported to-date.^19,20^

Protein engineering using directed evolution based approaches can be used to tweak substrate selectivity and drastically increase turn-over rates by orders of magnitude to facilitate commercialization of chemoenzymatic methods.^21^ There has been therefore interest in the last decade to develop new cell-based screening methods to identify novel specificities and improved catalytic performance of GSs and GTs to synthesize designer oligosaccharides, polysaccharides, and glycoconjugates.^21–29^ Directed evolution of GSs would also allow identification of beneficial mutations outside the active site that are nearly impossible to predict using rational approaches, particularly in the absence of any structural information. Additionally, extremophilic GHs offer an opportunity to develop novel GSs with higher specific activity that would favor glycan synthesis and improve reactant and/or substrate solubility under industrially-preferred bioprocessing conditions. However, one of the major challenges identified has been the lack of suitable ultrahigh-throughput screening (uHTS) methods for screening large GS and GT libraries (>10^6^ mutants/day). Withers first reported a two-plasmid uHTS method where one plasmid contained the GS gene while the other contained a GH-screening enzyme that only releases a fluorophore from the product of the GS reaction but not the original reactants.^30^ Similarly, chemical complementation using a yeast three-hybrid system was used to link GS activity to the transcription of a reporter gene, making cell growth dependent on GS reaction product formation.^22^ Both these approaches are highly specific to the individual GS family and have limited applicability to screen for novel GS specificity. The first universal HTS method to screen GS libraries (~10^4^ mutants/day) using glycosyl fluoride as the sugar donor was a pH based assay.^24^ Here, hydrofluoric acid, an end-product of the GS reaction using glycosyl fluorides, was detected by a pH sensitive colorimetric indicator. Also, a chemical probe that reacts specifically to the fluoride anion to generate a weak fluorophore was also used recently to screen small GS mutant libraries (~10^2^ mutants/day).^25^ However, to increase the probability of finding rarer GS mutants, fully-automated uHTS techniques capable of handling much larger mutant libraries (10^6^-10^13^ mutants/day) are necessary. Furthermore, other drawbacks with the current fluoride detection based HTS methods for GS engineering are: (i) low sensitivity limit for detection of reaction products (0.01-10 mM concentration range) which greatly reduces throughput and makes it challenging to fine-tune selection threshold, (ii) inability to distinguish between desired GS activity oligosaccharide products versus side-reaction products due to self-condensation of donor sugars and particularly due to hydrolysis of glycosyl fluorides due to its poor stability in aqueous conditions (e.g., glycosyl fluorides half-life stability ranges between few hours to days for most α- and β-anomers), and (iii) the lack of a sensitive fluorophore than can readily detect unreacted glycosyl fluoride reactant (or released fluoride product), which overall prevents the use of common cell sorting methods necessary to increase screening throughput.^18,31^ Withers and co-workers developed the first fluorescence-activated cell sorting (FACS) method for the directed evolution of sialyltransferases to screen for novel GTs with desired substrate specificity in a truly ultrahigh-throughput manner (>10^6^ mutants/day).^27^ FACS based uHTS methods alleviate the need to lyse cells, isolate plasmids, and retransform cells for iterative and fully-automated screening of much larger mutant libraries. Importantly, FACS based screening led to the discovery of a mutant GT with 400-fold increase in catalytic efficiency.^27^ Recently, Yang and co-workers have applied a similar FACS based screening approach to identify novel fucosyltransferase mutants with 6-14 fold increased catalytic efficiency (i.e., k_cat_/K_M_) for synthesis of fucosylated oligosaccharides like 3’-fucosyllactose.^32^ Similar directed evolution based experiments for GSs are necessary to identify novel mutants with increased catalytic efficiency and/or substrate specificity.^25,33^ The challenge is to use donor/acceptor substrates without directly incorporating fluorophore tags to monitor GS reactions, since that often results in biased mutants selection with fluorophore substrate specificity, highlighting a common issue with HTS assays – *you get what you screen for!*

Here, we report a novel fluorescence based uHTS approach for click-chemistry enabled detection of glycosyl azides as donor sugar reactants (versus released azide products) for rapid *in-vivo* detection of GS activity. This ultimately led to the development of a directed evolution methodology for screening GSs capable of synthesizing fucosylated glycans. The proposed universal uHTS assay can be used for the directed evolution of any class of GSs employing glycosyl azides as activated donor sugars. We showcase how this method was developed and applied to identify novel mutant GSs for the chemoenzymatic synthesis of several model fucosylated glycans. This method was developed based on a serendipitous discovery of distinct differences in relative fluorescence of the triazole-containing fluorophore product formed during the click-chemistry reaction of organic glycosyl azides versus inorganic azides. This difference in fluorescence intensity formed the basis for FACS-based sorting of *E. coli* cells expressing mutant GSs and thus containing different concentrations of GS reaction products depending on the mutant. Our validated uHTS method ultimately allowed rapid screening of a large library of fucosynthase variants (~10^6^ mutants) expressed in *E. coli* to identify several novel GS mutant sequences with increased activity for β-fucosyl-azide towards desired model acceptor sugars (e.g., pNP-xylose, lactose) to demonstrate several applications of this uHTS method.

## Results

### In-vitro click-chemistry detection of inorganic azide versus glycosyl azide

A red fluorophore dye tagged with dibenzocyclooctyne moiety, connected by a PEGylated linker, was a standard commercially available click-chemistry reagent (i.e., DBCO-PEG4-Fluor 545; see Supplementary Figure S1) that was reacted with either sodium azide (inorganic azide) or β-D-glucopyranosyl azide (organic azide) at 37 °C in 1X pH 7.4 PBS buffer for a total reaction time of 5 hours to continuously monitor the strain-promoted azide-alkyne cycloaddition (SPAAC) reaction kinetics (**Figure 1A**). The progress of the SPAAC reaction was quantitatively monitored by measuring the solution absorbance at 309 nm wavelength (λ_309_), which is the characteristic wavelength for alkyne groups.^34^ A decay in the measured absorbance at λ_309_ nm is indicative of the SPAAC reaction between the alkyne group in the dibenzocyclooctyne or DBCO moiety along with either the inorganic azide or organic azido groups (**Figure 1B**). Our results were consistent with Popik and co-workers who also reported a similar decrease in UV absorbance at 309 nm during the SPAAC reaction of 0.5 mM DBCO moiety with 2.5 mM sodium azide in PBS buffer (but in the presence of 5% methanol in that previous study).^34^ The SPAAC reaction kinetics data was fitted to a simple exponential decay function to obtain the apparent rate constants for the formation of the triazole moiety. The rate constants for sodium azide (0.0412+0.006 s^−1^) and glucosyl azide (0.043+0.003 s^−1^) were similar and therefore suggested that the SPAAC reaction proceeds at comparable rates for organic and inorganic azides.

**Figure 1.**
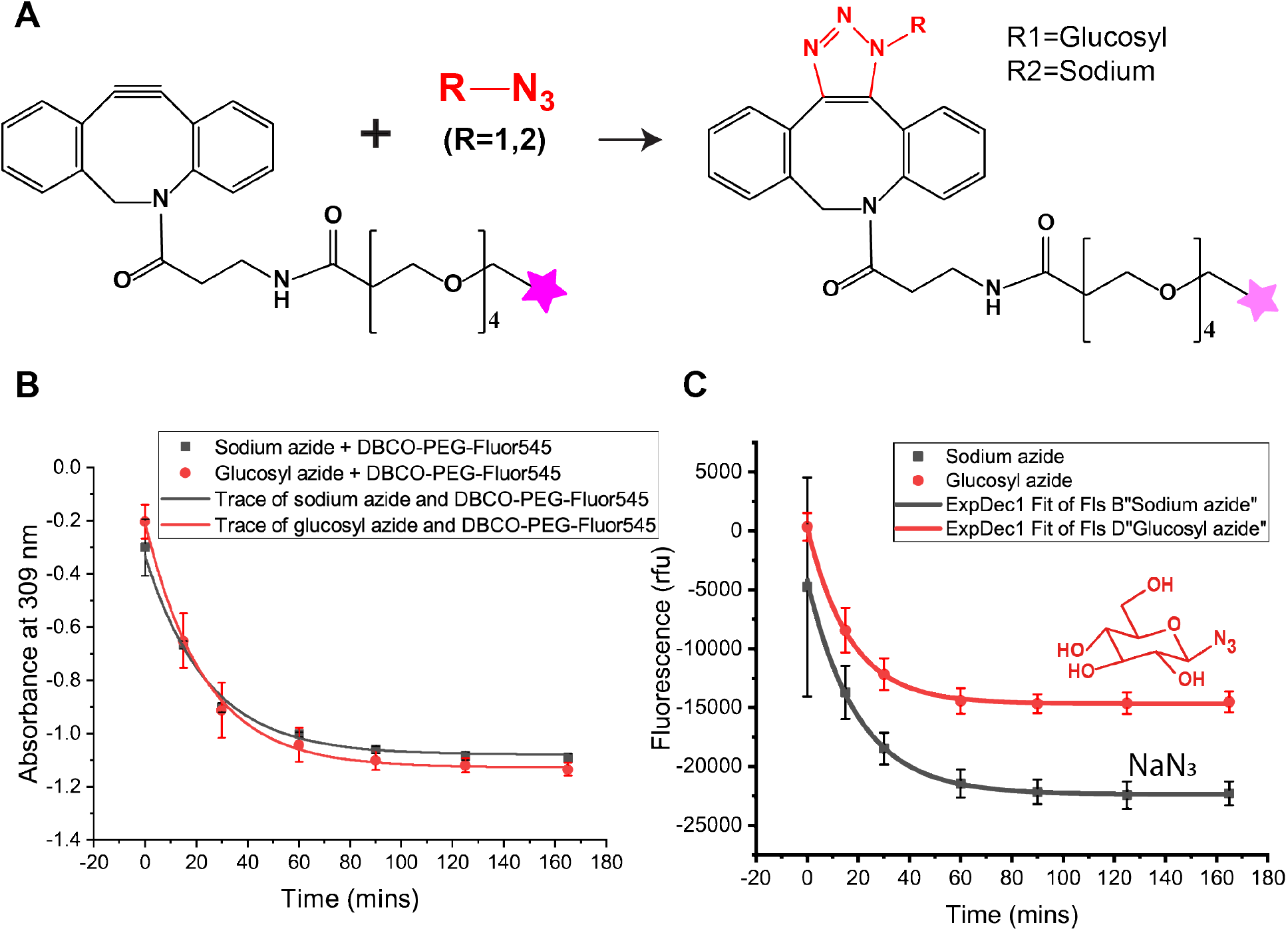
Triazole products of organic (β-D-glucopyranosyl azide) versus inorganic (sodium azide) azides formed during SPAAC reaction with DBCO-PEG4-Fluor545 exhibit different relative fluorescence intensities. A) Click chemistry or SPAAC reaction between DBCO-PEG4-Fluor 545 and azides result in a triazole product depending on the type of organic versus inorganic azide that reacts with the DBCO moiety. The relative change in observed triazole moiety specific absorbance (B) and overall product fluorescence (C) during the SPAAC reaction between DBCO-PEG4-Fluor 545 (200 μM) with either sodium azide or β-D-glucopyranosyl azide (400 μM) substrates in 1X PBS buffer, pH 7.4 at 37°C are shown here. Interestingly, while no difference in absorbance is seen, clear differences in final fluorescence readings can be seen for the SPAAC reaction products of organic versus inorganic azides. Note that Absorbance was measured at 309 nm, while, the fluorescence was measured at 550 nm excitation and 590 nm emission (with 570 nm cut off). Grey squares and red circles correspond to experimental data collected at noted time points for the inorganic and organic azide, respectively. The line traces are shown to aid the reader in following the relative change in values. Error bars shown here represent one standard deviation from the reported mean value.

Additionally, we had also monitored the solution fluorescence for the SPAAC reaction mixtures at every timepoint in tandem with the absorbance measurements. The excitation (550 nm) and emission (590 nm) wavelength filters used to quantify the red fluorescence measurements were specific to the Fluor545 fluorophore or standard tetramethylrhodamine (TAMRA) dye. Interestingly, the kinetic traces of the fluorescence data indicated a sharp decrease in the solution fluorescence over a period of 30 minutes. This decrease in fluorescence is concomitant with the reduction in λ_309_ nm absorbance seen during the SPAAC reaction, also corroborating that the reaction was complete within ~30-60 mins for both glucosyl azide and sodium azide. However, an unexpected observation was the ~30% difference in the absolute fluorescence intensities of the corresponding triazole products formed during the SPAAC reaction of DBCO-PEG4-Fluor 545 with sodium and glucosyl azide, respectively (**Figure 1C**). This absolute decrease in fluorescence intensity could be attributed to the currently poorly understood photophysical interactions of the TAMRA red dye fluorophore group to either the glycosylated versus nonglycosylated triazole moiety, formed during the SPAAC reaction, to observe differential quenching in observed red fluorescence. Furthermore, we see no significant impact of lower reaction temperatures on this phenomenon as well (see Supplementary Figure S2). Also, based on the full absorbance and excitation/emission fluorescence data, there seems to be a maximum decay in emission fluorescence for the SPAAC products, versus the unreacted DBCO-PEG4-Fluor 545 dye, only close to the 550 nm excitation wavelength for the Fluor545 moiety (Supplementary Figure S3).

In order to gain a more mechanistic understanding of this phenomenon, we first tested if the free organic versus inorganic azide had any impact on the red fluorescence of a structural homolog of TAMRA dye (e.g., Rhodamine-B) alone. However, we saw no significant difference in the absolute fluorescence for Rhodamine-B dye in the presence of glucosyl-azide versus sodium azide, suggesting that the triazole moiety formation is likely critical to observing any differences (see Supplementary Figure S4 and Supplementary Text S1). Next, we tested if inter-molecular interactions of a glycosylated versus non-glycosylated triazole moiety with the Rhodamine-B dye had any impact on dye fluorescence. To test this hypothesis, the SPAAC reaction between a model DBCO-moiety lacking a fluorophore group (i.e., DBCO-NHS) and the azide substrates was first performed to produce a triazole product before addition of Rhodamine-B dye to the reaction solution. Interestingly, the addition of Rhodamine-B to either the glycosylated versus non-glycosylated triazole SPAAC products showed no significant change in fluorescence compared to the Rhodamine-B dye added along with DBCO-NHS by itself (see Supplementary Figure S4). This result suggests that intra-molecular interactions of the red fluorophore group with the triazole moiety, facilitated by the connected PEGylated linker, is necessary for the appropriate photophysical interactions that result in differential fluorescence observed for glycosylated versus non-glycosylated triazole-fluorophore based SPAAC products. The PEG linker likely facilitates intramolecular interactions between the triazole moiety, after reaction of DBCO moiety with either the azido sugar or free azide, and the FLUOR-545 group in well-defined molecular orientations to facilitate donor–acceptor interactions through intramolecular charge transfer (ICT) or fluorescence resonance energy transfer (FRET) type photophysical interactions. These interactions, though currently not well-understood, clearly result in a differential reduction in fluorescence for the SPAAC reaction products studied here. We also observed that changing the glycosyl moiety from glucose to fucose also did not alter the relative trends in fluorescence patterns noted here (see Supplementary Figure S5), suggesting that glycosylated triazoles likely behave similarly but more work is needed to investigate this phenomenon more closely for structurally distinct azido sugar isomers. Next, we took advantage of this serendipitous discovery to quantitatively estimate if this method could detect the extreme and intermediate varying concentration limits (e.g., unreacted glycosyl-azide substrate versus free released azide products) expected during a typical GS reaction (see Supplementary Figure 6), and subsequently developed a fluorescence based uHTS method for sorting cells expressing GSs based on these preliminary *in-vitro* results.

### TmAfc fucosidase is a potential glycosynthase but with poor fucosynthase activity

To further develop the uHTS technique, a α-L-fucosidase enzyme (Tm0306 gene; Genbank Accession ID NC_000853) isolated from *Thermotoga maritima*,^35^ was chosen as a model GS enzyme for bacterial expression and further *in-vitro* testing (**Figure 2A**). This fucosidase (TmAfc0306) has been engineered to an α-L-fucosynthase by mutating the catalytic nucleophile D224 residue to either alanine, glycine, or serine mutants (Supplementary Figure S7).^31^ Our chemical rescue experiments on the mutants revealed that an exogenous nucleophile, azide was able to rescue the hydrolytic activity of D224G by 98% while alanine and serine did not show any significant recovery in activity (Supplementary Figure S8). The *in-vitro* reaction of pNP-β-D-xylose (acceptor sugar) and β-L-fucosyl azide (donor sugar) with purified D224G mutant resulted in the formation of only two minor glycosynthase products α-L-Fuc-(1,4)-β-D-Xyl-pNP (55%; molar basis) and α-L-Fuc-(1,3)-β-D-Xyl-pNP (45%; molar basis) (**Figure 2B** and Supplementary Figure S9). The mechanism of the fucosynthase reaction is outlined in **Figure 2C** where the acid/base residue of D224G mutant deprotonates the −OH group at the C4 position of pNP-β-D-xylopyranoside. Next, the deprotonated pNP-β-D-xylopyranoside anion would attack the anomeric carbon of β-L-fucopyranosyl azide present adjacent to the catalytic nucleophile D224G site facilitating the release of the azide leaving group. Finally, pNP-β-D-xylopyranoside would form a glycosidic bond with β-L-fucosyl moiety to produce β-L-fucopyranoside-β-D xylopyranoside-pNP as the final GS reaction product. However, the total GS reaction product yield was found to be only ~6% (i.e., based on initial pNP-β-D-xylose starting concentration), even after prolonged reaction incubation period of several days, indicating that the D224G GS activity is very low. Hence, D224G was used as the baseline GS to identify additional mutants using our uHTS method to enhance its native fucosynthase activity.

**Figure 2.**
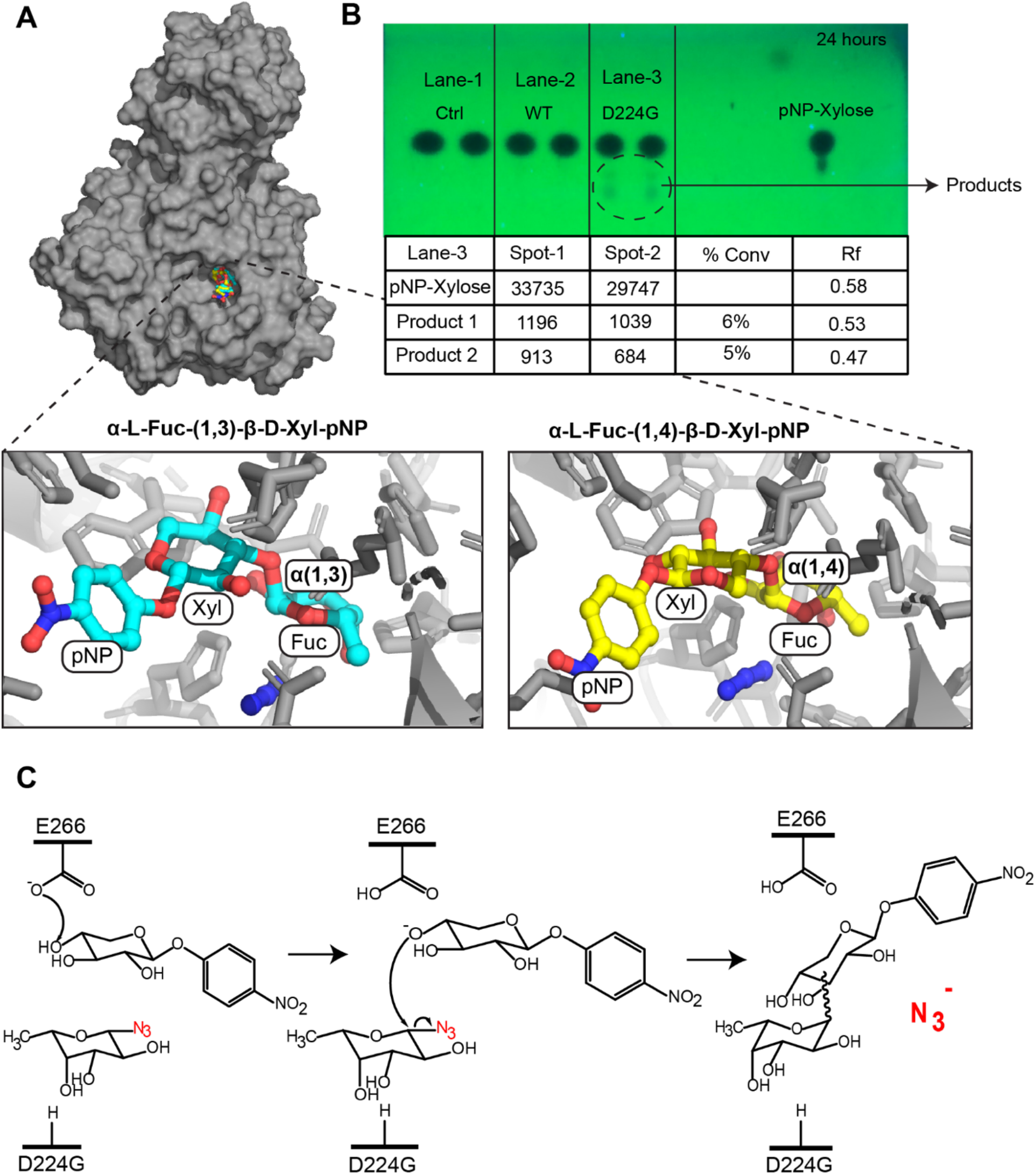
TmAfc-D224G is α(1,3) and α(1,4) specific for glycoside synthesis with pNP-β-D-xylose and β-L-fucopyranosyl azide. a) Glycosynthase products docked in the active site of TmAfc-D224G based on the quantum mechanical/molecular mechanics^36^ predicted structures shown here. b) UV image of the TLC analysis of glycosynthase reaction of D224G with β-L-fucopyranosyl azide and pNP-β-D-xylopyranoside in 50 mM MES pH=6.0 at 60°C. Here, product 2 is α-L-Fuc-(1,3)-β-D-Xyl-pNP and shown on left/cyan, while product 1 is α-L-Fuc-(1,4)-β-D-Xyl-pNP and shown on right/yellow. c) Reaction scheme of glycosynthase reaction of Tm0306_D224G with pNP-β-D-xylose and β-L-fucopyranosyl azide.

### Conceptual overview of the uHTS methodology

The overall approach for screening glycosynthases using SPAAC or click-reaction based *in-vivo* detection of GS activity and FACS based cell sorting is outlined in **Figure 3**. This approach was used to screen mutants of D224G with improved fucosynthase activity using β-L-fucosyl azide as the donor and pNP-β-D-xylose as the acceptor. The parent/template DNA (i.e., D224G) was first diversified to create a random mutagenesis library using the standard error-prone Polymerase Chain Reaction (epPCR) methodology (step-1). The average number of mutations introduced during epPCR were estimated to be ~3-4 per mutant construct. The epPCR mutant library obtained was cloned into a T5-promoter based plasmid,^37^ and then transformed into standard *E. coli* (E.cloni) competent cells in step-2. The transformed cells were grown overnight and then diluted down by 1:20 for further incubation at 37 °C until the cells reached mid-exponential growth phase (i.e., OD_600_ between 0.4-0.6). In step-3, protein expression was induced using IPTG for 1 hour at 37 °C. After protein expression was verified (i.e., using chemical rescue activity assay for crude cell lysates), in step 4, the glycosynthase reaction was carried out *in-vivo* for 2 hours at 37 °C using freshly-added GS reaction substrates pNP-β-D-xylose and β-L-fucopyranosyl azide (indicated in **Figure 3** as orange-yellow and red-blue colored symbols, respectively). Step-4 schematic also outlines the two extreme possibilities of the GS reaction outcomes expected for the mutant library. In one scenario, there is no GS reaction taking place and thus the cells ideally should contain only unreacted azido sugars. In the second scenario, assuming the GS reaction goes to 100% completion, then the cells would contain only the released free azide ion product (indicated in **Figure 3** as a blue circle). At the end of the GS reaction completion in step-4, DBCO-PEG4-Fluor545 was added to the cells to trigger *in-vivo* click reaction at 37 °C for at least 30 minutes as part of Step-5.

**Figure 3.**
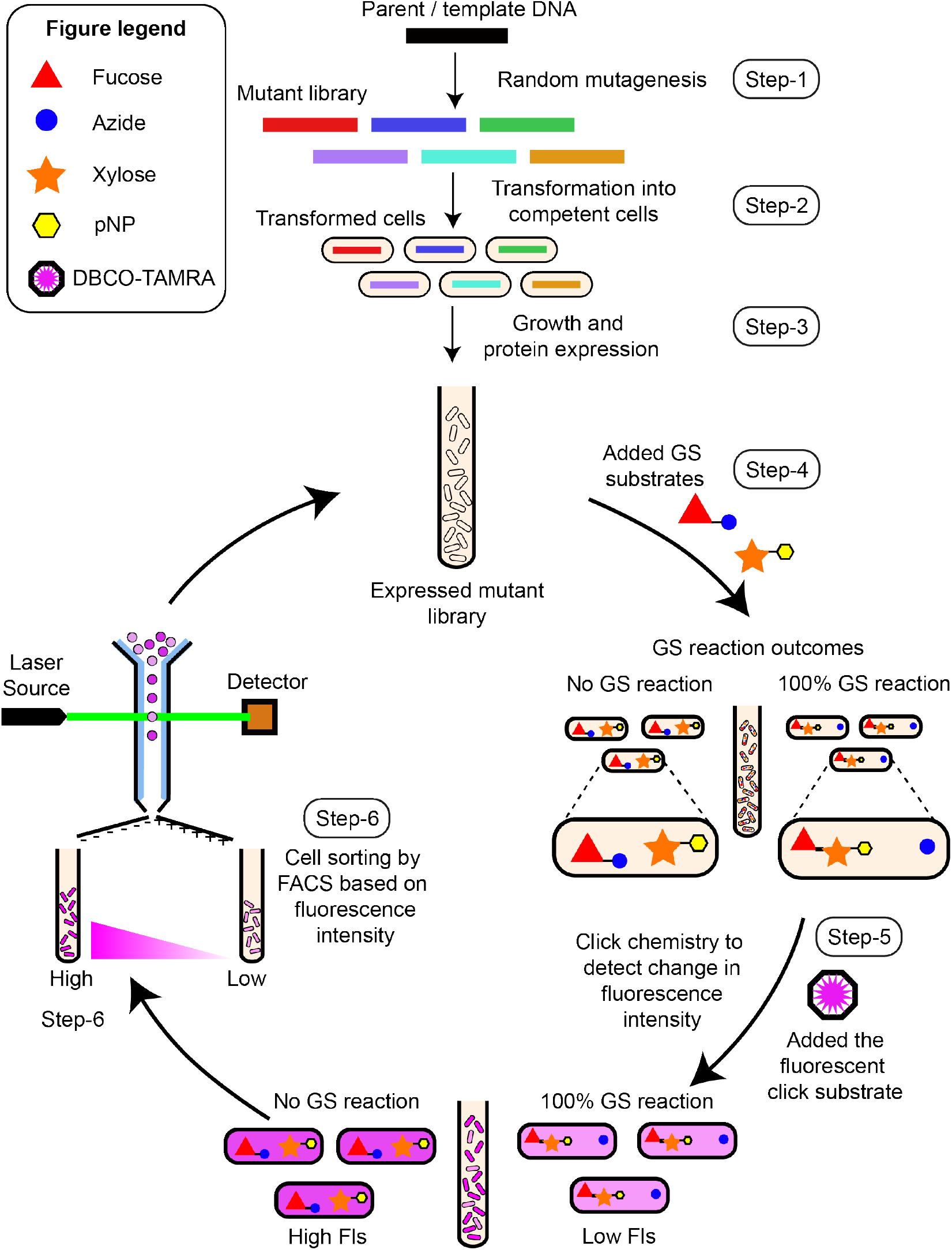
Conceptual overview of click-chemistry based ultrahigh-throughput screening (uHTS) method for in-vivo directed evolution of glycosynthases. Here, we first generate a mutant library (step 1), introduce the library into expression strains (step 2), and then grow cells to desired OD before expression glycosynthase mutants (step 3). Next, we add the substrates for performing the glycosynthase reaction (step 4) followed by adding the SPAAC reagent (DBCO-TAMRA) for click-labeling and detection of cells containing lower concentration of azido sugars (step 5). Cells with lower fluorescence intensity (FI) are then sorted using FACS (step 6) prior to isolating novel glycosynthase mutants or repeating another round of uHTS using a directed evolution approach to enrich mutants sorted in previous rounds of screening.

As seen for both the *in-vitro* and *in-vivo* SPAAC reaction results (**Figure 2B** and Supplementary Figures S6 and S10), a decrease in SPAAC reaction products fluorescence was seen for free azide versus unreacted azido sugars alone. For example, if the GS reaction went to completion then the fluorescence decrease in scenario 2 is expected to be higher due to the presence of higher relative concentration of free azides compared to the lesser reduction of fluorescence in scenario 1 if no GS reaction takes place (i.e., higher relative concentration of azido sugars). The relative difference in fluorescence intensities forms the fundamental basis for single-cell separation using FACS (in step-6). The cell population with lower versus higher fluorescence intensities were sorted into separate tubes containing recovery/growth media. The recovered cells were then subjected to a second round of FACS by repeating steps 3-6. During the second round of sorting, individual cells with lower fluorescence were collected in a 96-well plate for further characterization of the isolated GS mutants. Mutants identified after the second round of cell sorting with lower fluorescence intensity indicate that multiple rounds of library sorting can result in enrichment of highly active GS mutants. The cells isolated from the second sort in 96-well plates were next tested for their crude cell lysate based GS activity using the chemical rescue colorimetric assay method with pNP-fucose as substrate in the presence of exogenous azide nucleophile. Suitable controls were also included for the FACS screening to account for any false positives. Isolated mutants from the second round of cell sorting resulted in chemical rescue with higher activity compared to the template control (i.e., D224G) and were identified as positive mutants. The positive mutant clones were finally isolated for further characterization (e.g., DNA sequencing, expression, and in-vitro GS assays).

### Isolation of novel TmAfc mutants with improved fucosynthase activity

Before applying the uHTS approach outlined above to facilitate *in vivo* sorting of TmAfc-D224G epPCR mutant library, we ran several *in-vitro* and *in-vivo* control experiments as mostly discussed in the supplementary information document. Briefly, we first tested the sensitivity of *in-vitro* SPAAC reaction in the presence of various mixtures of inorganic and organic azides to provide proof-of-concept results that total SPAAC reaction solution fluorescence is a function of reactants composition. For example, an equimolar mixture of sodium azide and β-D-glucopyransoyl azide reacted with DBCO-PEG4-FLUOR 545 gave a fluorescence intensity that was in the intermediate range of intensities observed for sodium azide and β-D-glucopyranosyl azide alone (Supplementary Figure S6). These *in-vitro* results suggest the applicability of the SPAAC reaction to detect mixed concentrations of unreacted sugar azide and released inorganic azide formed by active GSs of varying degrees of catalytic efficiency. Therefore, in principle, the SPAAC reaction using DBCO-PEG4-FLUOR 545 should be also able to give a differential fluorescence response for GS products formed and/or unreacted substrates present under *in-vivo* conditions as well. In order to validate this, confocal fluorescence microscopy was first performed to confirm that the fluorescent SPAAC reagent (i.e., DBCO-PEG4-FLUOR 545) could readily permeate inside *E. coli* cells (Supplementary Figure S11 and Supplementary Text S2). Flow cytometry also confirmed efficient permeation of the SPAAC reagents into the cells. Next, we used flow cytometry to determine total fluorescence intensity and distribution for *E. coli* cells containing SPAAC reaction products for DBCO-PEG4-FLUOR 545 reacted with either type of azide alone (Supplementary Figure S10). As seen during *in-vitro* assays, *E. coli* cells containing SPAAC reaction products for glucosyl-azide clearly gave a different fluorescence intensity than the cell population containing SPAAC reaction products for sodium azide alone. We next used flow cytometry (and FACS) to confirm if *E. coli* cells expressing D224G control fucosynthase gave a clearly distinguishable decrease in fluorescence intensity compared to TmAfc wild-type GH control after conducting GS and SPAAC reactions using similar steps as outlined in **Figure 3** (and Supplementary Figure S12). Overall, these proof-of-concept uHTS methodology development results all suggested it would be possible to sort mutant GSs with increased activity based on decreased fluorescence expected for the SPAAC reaction products.

In order to test our uHTS methodology, we performed random mutagenesis using TmAfc-D224G as a template (control) and sorted the epPCR mutant library in a BD Influx High Speed Sorter to identify mutants with increased fucosynthase activity towards pNP-xylose as acceptor after two rounds of FACS sorting. See Supplementary Text S3 and Supplementary Table S2 for additional details on the epPCR method. The fluorescence intensities of unstained *E. coli* cells (negative control) and cells expressing template D224G protein (positive control) were first captured to optimize the FACS instrument parameters (e.g., pressure, gain) and build the fluorescence gates for sorting (**Figure 4A**). Two distinct populations with fluorescence intensities with range differing over several orders of magnitude were clearly observed when the epPCR mutant library cells were analyzed using FACS. Two fluorescence gates, differing over a minimum 10-fold magnitude, called ‘Low’ and ‘High’ were drawn to identify and classify these cell populations (Supplementary Figure S13). The cells which had fluorescence in the ‘Low’ gate were separated by FACS and collected into a single tube containing LB recovery/growth media. The sorted cells were regrown and subjected to second round of FACS sorting to eliminate false positives. During the second round of sorting, individual cells in the ‘Low’ gate were sorted and collected as individual cells in a 96-well plate with LB recovery/growth media.

**Figure 4.**
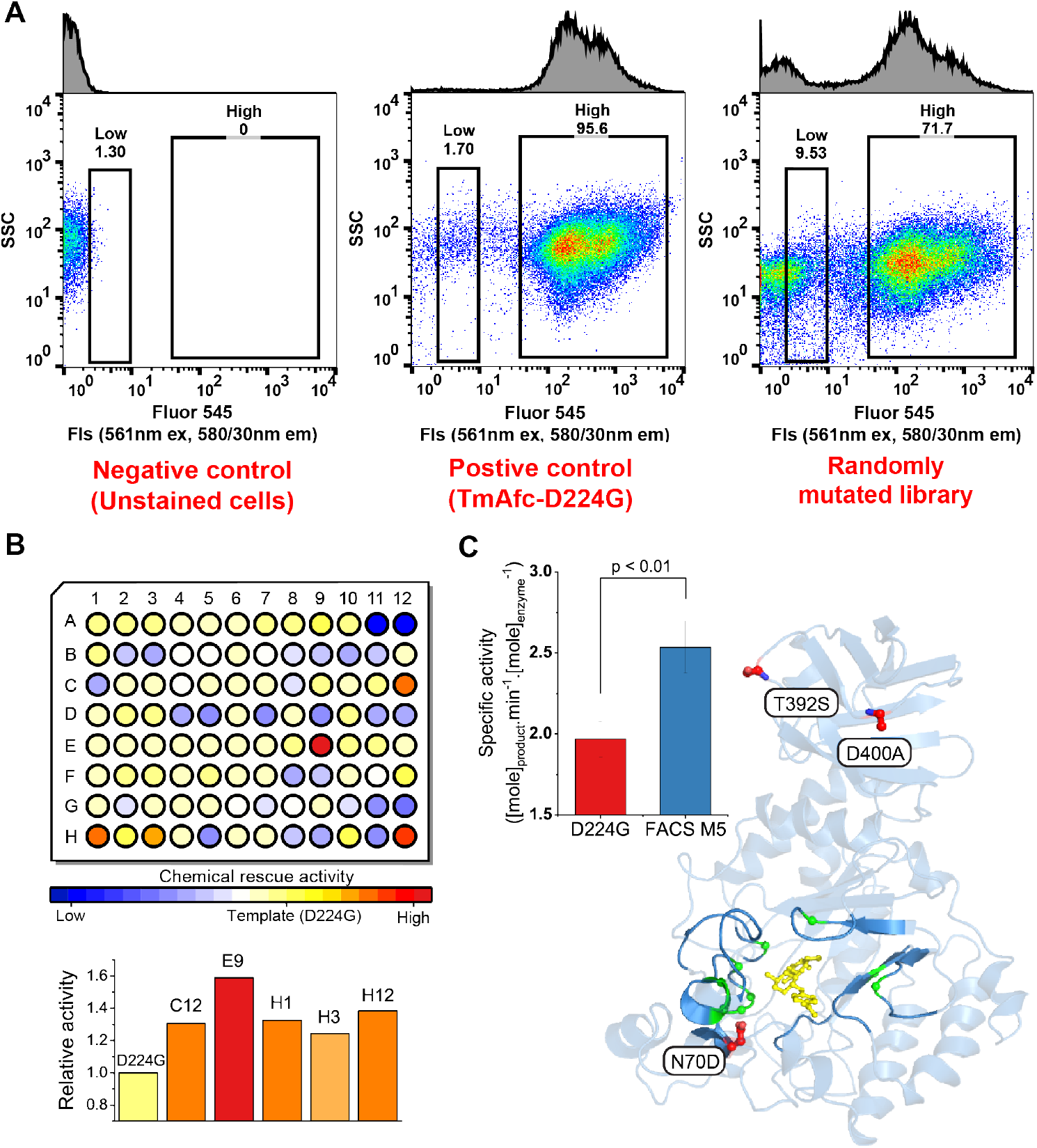
Improved fucosynthase mutants identified, compared to starting template (TmAfc-D224G), by proposed uHTS method. A) Scatter plots (SSC vs Fls 561nm ex/580/30nm em) for unstained cells, cells with D224G template, and randomly mutated library are shown here. The gates “Low” and “High” are indicated as rectangles. The cell population percentage in each gate is indicated by the number above the rectangle. B) Chemical rescue activity heatmap for cells sorted into 96-well plate after second uHTS round is shown. Here, A1 to A10 are the D224G controls and while all other wells are FACS sorted variants. Data for improved mutants is shown as orange to red color (normalized to D224 control data set to 1.0). The mutants/cells which showed no rescue of activity or did not grow after sorting are indicated by blue color. This screening step eliminated any false positives and only mutants with higher chemical rescue activity compared to TmAfc-D224G were chosen for DNA sequencing. C) FACS M5 variant (also H1 location on 96-well plate) sorted from the epPCR library was expressed, purified, and tested to show a 29% improved overall fucosynthase specific activity compared to the D224G control. The specific activities of both the TmAfc-D224G and FACS M5 mutant were determined from the initial rates of reaction between β-L-fucopyranosyl azide and pNP-β-D-xylopyranoside to form glycosynthase products (i.e., α-L-Fuc-(1,3)-β-D-Xyl-pNP and α-L-Fuc-(1,4)-β-D-Xyl-pNP). DNA sequencing revealed three mutational sites in the M5 construct (shown in red; N70D, D400A, T392S). The residues in the active site of the M5 mutant enzyme interacting with the docked substrates are shown as green spheres. Error bars shown here represent one standard deviation from the reported mean value.

The cells were then grown, and protein expression was induced in a 96-well culture plate. To these induced cell cultures, the cells were lysed and the substrate pNP-Fucose along with the external nucleophile sodium azide was added to check for expressed enzyme chemical rescue activity as a secondary screen prior to DNA sequencing. The FACS sorted single-cell epPCR mutants with improved chemical rescue activity compared to the template D224G control, as shown in **Figure 4B**, were then selected for plasmid DNA extraction and subsequent DNA sequencing of identified positive GS mutants. Unique mutants identified after DNA sequencing were individually expressed and purified proteins were used to perform *in-vitro* glycosynthase reactions. From our current analysis, we have identified at least four unique fucosynthase mutants that give much higher glycosynthase activity (**Figure 4B-C** and Supplementary Text S4). Interestingly, N70D was the highly conserved mutation identified for all four mutants, which in combination with T392S and D224G mutations gave significantly higher fucosynthase activity compared to the original template (D224G). One of these mutants (FACS M5) carrying three new mutations, in addition to D224G (or TmAfc-D224G-N70D-T392S-D400A), was expressed and purified to conduct systematic in-vitro enzyme activity assays (Supplementary Figure S7). We have further confirmed the identity of both the GS reaction products that were first qualitatively characterized using TLC and then quantitatively analyzed using HPLC-UV analysis. The specific activities of the M5 mutant, determined from the initial rate of the glycosynthase reaction between β-L-fucopyranosyl azide and pNP-β-D-xylopyranoside, was 29% greater than the template D224G (**Figure 4C**; see inset bar graph). Here, wild-type fucosidase (with intact nucleophile at D224) gave no measurable glycosynthase activity and therefore no fucosynthase activity data is not shown here for this construct. Mutation of the asparagine residue (N70D) was the most highly conserved residue near the active site for nearly all identified mutants with improved fucosynthase activity. It is likely that N70D mutation could alter the interaction of neighboring Trp residue and helps with improved docking of the substrate in the active site (**Figure 4C**; in yellow). However, more detailed computational and experimental analysis is required to explain why some of the other distal site mutations result in improved fucosynthase activity compared to the original D224G template.

### Computational modeling of the M5 mutant to rationalize improved activity

The M5 mutant construct was modeled and simulated to investigate structural features behind the improved glycosynthetic activity compared to the D224G single mutant. Unbiased molecular mechanics (MM) simulations were used to characterize the structural and dynamic changes associated with the mutations (**Figure 5A**) and hybrid quantum mechanics/molecular mechanics (QM/MM) simulations were used to obtain the free energy profile of the α(1,4) glycosynthetic reaction within the enzyme active site (**Figure 5B**).

**Figure 5.**
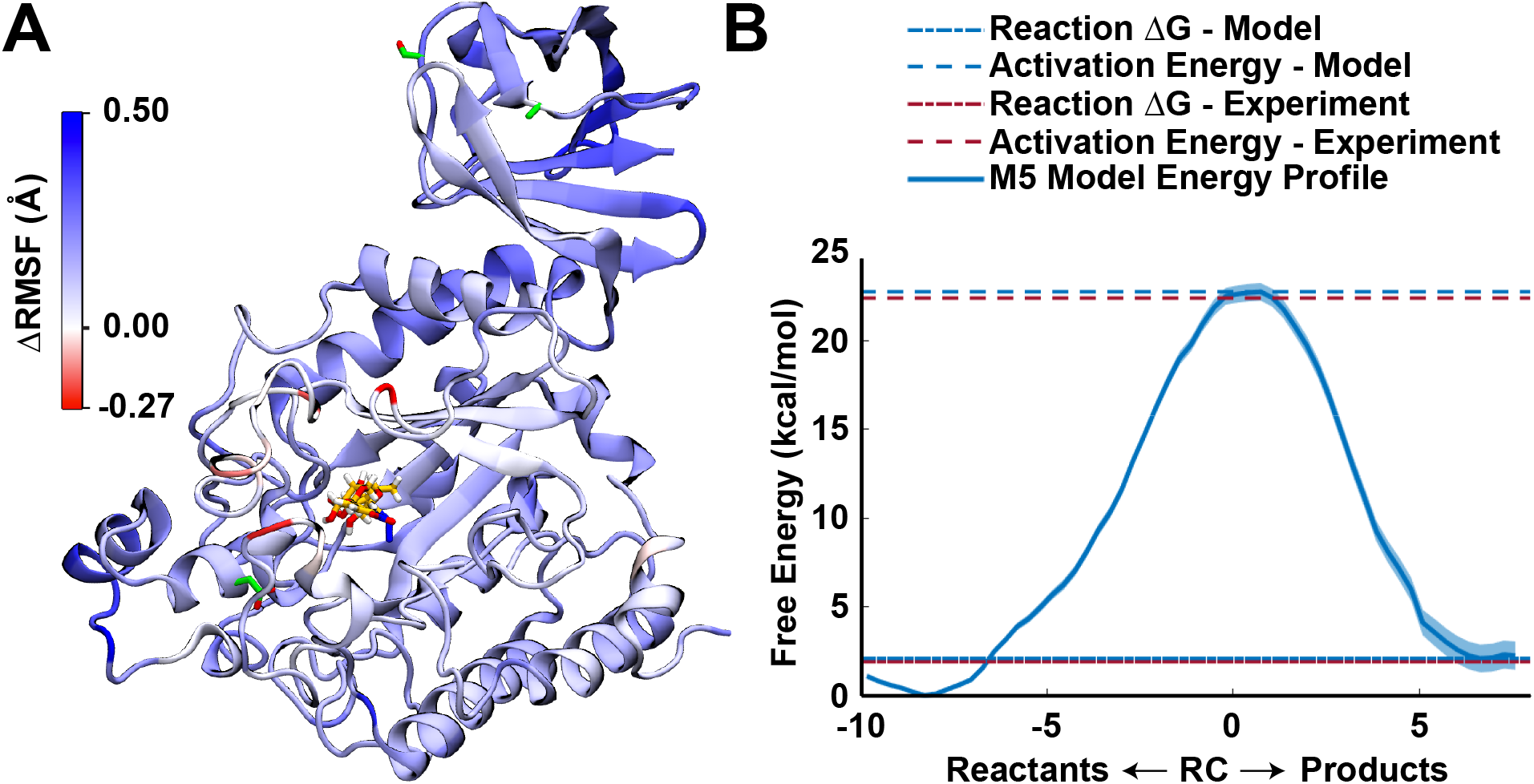
Root mean square fluctuation of M5 mutant fucosynthase structural model and QM/MM simulations predicted free energy profile of fucosynthase reaction. A) Snapshot of the enzyme where each residue is colored according to the change in root mean square fluctuation (ΔRMSF) associated with the three additional mutations in the M5 construct compared to the D224G mutant (new mutations shown in green). An increase in RMSF (blue colors) indicates that the mutations resulting in a destabilization of that residue, while a decrease (red colors) indicates stabilization. The substrates in the binding pocket are shown in yellow. B) The free energy profile from the QM/MM umbrella sampling simulations for the M5 construct α1,4 reaction. The error bars represent uncertainty in the process of fitting the profile using the multistate Bennett acceptance ratio. “Activation energy” refers to the forward (reactants to products) energy barrier while “reaction ΔG” refers to the difference in free energy between the product and reactant states. The red dashed lines are taken from the experimental results reported in this study, based on the initial reaction rate (for the activation energy) and the equilibrium ratio of products to reactants (for the reaction ΔG), respectively. The blue dashed lines are the corresponding values taken from the QM/MM model free energy profile, for comparison.

The three additional mutations in the M5 construct did not greatly change the protein conformation, but did result in a decrease in the rigidity of almost all parts of the enzyme (**Figure 5A**). This result suggests that the means by which the M5 mutant improves upon the glycosynthetic activity of the single-mutant is by loosening the highly specific structure of the wild-type active site (which would have evolved to suit glycoside hydrolysis exactly) in favor of a more general α-fucosyl oligosaccharide binding site with a de-emphasized preference for hydrolysis over synthesis. As previous authors have pointed out, de-specialization is the expected outcome of early rounds of directed evolution, as the enzyme first “unlearns” its specificity for the wild-type reaction before it can begin to develop new specificity for the target reaction.^38,39^

The QM/MM simulations were in extremely close agreement with the experimental results, both in terms of activation energy and overall reaction ΔG (**Figure 5B**). This close agreement strongly suggests that the reaction step is rate-limiting in turnover of this enzyme, in which case the mechanism of increased activity in the M5 construct over the single mutant would be a reduction in the forward reaction activation energy barrier, albeit one too small to confidently distinguish with this computational model (based on the experimental results, the activation energy should change by just 0.15 kcal/mol at ~333 K). Taken in the context of the MM results, the most likely explanation for the improved activity is a subtle loosening in the tightness of the active site in such a way as to better permit the glycosynthetic transition state, perhaps for example by making additional room for the slightly longer C-N bond in the glycosynthase over the shorter C-O bond in the wild-type enzyme.

## Discussion

Azido sugars have been used extensively as bio-orthogonal reagents in the last two decades to unravel the inner-workings of the glycosylation pathways of cellular systems both at the singlecell and organismal levels.^40,41^ However, azido sugars or glycosyl azides have not yet been used extensively as a reagent for synthesis of glycans. Nevertheless, these reagents offer several advantages over other activated sugar donors like glycosyl fluorides or nucleotide sugars for *in-vitro* chemoenzymatic synthesis of glycans. Glycosyl azides can be readily chemically synthesized using one-pot reactions from unprotected sugar monomers, as well as produced enzymatically at high yields, unlike other activated donor sugars like glucosyl fluorides.^31,42,43^ Furthermore, GHs are being rapidly discovered through cheaper sequencing of diverse microbial, microbiome, and metagenomic sources.^13,44^ These GHs offer a large selection of enzymes that have not yet been exploited for engineering more effective and highly selective GSs for desired donor sugars. Directed evolution of GSs can be used to increase reaction rate and introduce novel donor sugar or acceptor group substrate specificity.^21^ However, currently there are very limited options available for universal HTS methods for rational engineering or cell-based directed evolution of GSs.^24,25,30,45^ Specifically, other than this report, there are currently no HTS methods available that facilitate ultrahigh-throughput cell-based screening and directed evolution of GSs using azido sugars as glycosyl donors.^18^ Here, we have now developed, validated, and applied a novel SPAAC or click-chemistry enabled uHTS method for screening large GSs libraries. As proof of concept, we showcased how our screening technique facilitates rapid fluorescence activated cell sorting of a large library of glycosynthase variants (>10^6^ mutants) expressed in *E. coli* to identify several novel mutants with increased activity for β-fucosyl-azide activated donor sugars towards desired acceptor sugars, demonstrating the broader applicability of this methodology.

One of the major advantages of using glycosyl azides as substrates for GS reactions is that the azido moiety can be selectively conjugated to alkyne-based fluorophore groups using a modified Staudinger ligation or copper-free click-chemistry under reaction conditions compatible with the *in vivo* environment.^40,46–48^ Similar approaches can be utilized for cell-free based ultrahigh-throughput screening methods for CAZyme engineering as well.^49^ Interestingly, the relative difference in the fluorescence intensity of the glycosylated versus non-glycosylated triazole product influences the observed fluorescence for cells enriched in the corresponding SPAAC reaction products. Recent studies have suggested that the alkyne-derived substituent moiety of a “click” triazole can engage in electronic conjugation with the triazolyl core that may have profound influences on the optical properties of these compounds.^50^ Photon reabsorption behavior of such compounds can be vexing for the development of energy efficient light-emitting materials, however, this could be advantageous for producing a composite emission color that facilitates selective detection of differentially substitutes triazole products during SPAAC reactions. The influence of the triazole moiety substitution patterns on the electrochemical and photophysical properties of the donor-acceptor groups has been reported in the literature.^51^ However, there are no reports, to the best of our knowledge, on the effect of sugar-substituted triazole groups on the variable fluorescence quenching ability due to likely intramolecular photophysical or FRET type interactions with a red-fluorophore group like TAMRA or Rhodamine-B. Future work must resolve the mechanistic basis for selective reduction in fluorescence of non-glycosylated triazole products to facilitate design of more efficient SPAAC reagents to reduce false positives detection using a similar uHTS approach.

Moracci and co-workers had shown how sugar azides can be used for synthesis of glycans using thermophilic GH29 fucosidase enzyme that are able to accommodate slightly larger leaving groups like azides in the mutated D224G active site, instead of smaller leaving groups like fluorides.^31^ However, while the D224G was reported to show some fucosynthase activity, we found that the overall activity of this enzyme is actually quite low. Mayes and Burgin recently investigated the first unbiased transition path sampling study of this GH29 fucosynthase enzyme to reveal a single-step mechanism with an oxocarbenium-like transition state.^36^ Their study suggested one potential explanation for the poor reaction efficiency observed for the D224G fucosynthase was that there were no nearby residues or water molecules to stabilize the departure of the azide group. This could explain why the D224S mutant gave an inactive fucosynthase unlike the D224G due to the relatively bulky serine side chain that likely further restricts azide group accessibility. We have now successfully isolated several novel fucosynthase mutants with improved activity for β-fucosyl-azide towards pNP-xylose. Interestingly, we have identified N70D as a mutation close to the active site that we hypothesize plays an important role in stabilizing substrate binding. However, distal mutations identified outside the active site could not be easily predicted rationally. Some of these distal site mutations may participate in stabilizing and/or destabilizing parts of the protein fold that could help increase fucosynthase activity, as shown in **Figure 5A**. The two major glycosynthase products, α-L-Fuc-(1,4)-β-D-Xyl-pNP and α-L-Fuc-(1,3)-β-D-Xyl-pNP, were also formed for the improved mutants as well but at much higher turnover rates than previously seen for the D224G variant. We suspect the additional mutations identified during uHTS screening could have further stabilized the intermediate oxocarbenium-like transition state by aiding in (or equivalently, resisting less) the expulsion of the azide leaving group to increase GS catalytic efficiency for the formation of both α-1,3 and α-1,4 isomers. Interestingly, some of the mutations identified on M5 construct shifted the reaction equilibrium towards the α-1,3 versus α-1,4 isomers, suggesting that the uHTS methodology could be useful to also fine-tune glycosidic bond stereochemistry as well. For example, the D224G mutant also gave a ~55:45% (molar basis) of α-L-Fuc-(1,4)-β-D-Xyl-pNP:α-L-Fuc-(1,3)-β-D-Xyl-pNP, as also reported previously.^31^ However, the M5 mutant gave a glycosynthase product distribution of ~48:52% (molar basis) in favor of the α-1,3 isomer. Closer inspect of the *in-silico* mutated M5 structure revealed a native tryptophan residue (W67) adjacent to the N70D mutation that plays an important role in stabilizing the donor sugar and acceptor sugar via hydrogen-bonding and CH-π stacking interactions, respectively. The residues in the active site of the M5 mutant interacting with the docked substrates are shown in green (**Figure 4C**). Additional fucosynthase mutants with orders of magnitude higher glycosynthase activity than the reported M5 mutant are expected as one further optimizes the FACS ‘Low’ gate for capturing and isolating novel mutants with even lower fluorescence (NOTE: see <10^0^ fluorescence intensity scatter plot data in red/green contour displayed outside the ‘Low’ gate for epPCR library as seen in **Figure 4A** which suggests likely presence of novel GS mutants with even higher fucosynthase activity than M5), as well as screening additional mutant libraries generated using more advanced mutagenesis methods than epPCR.

Finally, this uHTS strategy is currently being exploited to sort mutant D224G epPCR library to identify novel GSs with altered substrate specificity by changing the acceptor sugar from pNP-xylose to either lactose, N-acetylglucosamine, or galactose (Supplementary Figure S14). Here, we can see a nearly 10-fold increase in the relative percentage of mutant cells identified in the low gate (~17% of total population) compared to the starting D224G control shown in **Figure 4** (1.7% of total population). These preliminary results clearly highlight that our proposed screening method could be easily applied to readily evolve and screen GS activity for novel acceptor sugars as well. Fucosylated glycans play a critical role in the selective recognition and metabolism of probiotic gut microbes. Human milk oligosaccharides (HMOs) are composed of a tetrasaccharide backbone that is selectively fucosylated to N-acetylglucosamine and/or galactose mirroring the four Lewis blood group antigens. All fucosyltransferases responsible for HMOs fucosylation are yet to be identified, which limits our knowledge on biosynthesis of HMOs. In particular, D224G showed low GS activity with fucosyl azide and lactose as substrates (data not shown). Furthermore, there have been significant advancements recently in the identification and sequencing of entire gene clusters that control the expression of GHs by probiotic gut bacteria (like *Bifidobacterium longum*) dedicated to bacterial growth on HMOs alone.^52^ One of the other major roadblocks remains the limited availability of complex glycan reagents that mimic the 100+ naturally occurring HMO to better understand the underlying mechanisms of action on human health. Identification of evolved and novel GS mutants capable of synthesizing 2’-fucosyl lactose, an essential HMO, along with several other Lewis blood group antigen oligosaccharides is currently under investigation in our laboratory. Future research will therefore focus on the development of novel glycosynthases that are selectively engineered and evolved using our uHTS method to identify novel mutants for efficient synthesis of complex HMOs or Lewis blood group antigens.

## Supporting Information

Supporting Information files are provided online as supplementary methods, supplementary results (as SI Figures S1-S15, Tables S1-S2, Text S1-S4), and supplementary discussion.

## Acknowledgements

SPSC acknowledges support from the US National Science Foundation CBET (Award No. 1704679 and 1904890) and Rutgers School of Engineering. This work used the Extreme Science and Engineering Discovery Environment (XSEDE),^53^ which is supported by National Science Foundation grant number ACI-1548562. Lastly, we are very grateful to other members from the Chundawat research group for their critical feedback and contributions during the course of this project.

## Data Availability

Authors confirm that all relevant data are included in the paper and/or supplementary information files.

