## Supplementary Information for "Click-chemistry enabled directed evolution of glycosynthases for bespoke glycans synthesis"

**For**

### Supplementary Information (SI) Table of Contents

**Supplementary Figure S1.** Molecular structure of DBCO-PEG4-Fluor 545. (Pg:3)

**Supplementary Figure S2.** *In-vitro* SPAAC chemistry reaction rates at varying temperature. (Pg:4)

**Supplementary Figure S3.** SPAAC emission fluorescence at varying excitation wavelengths. (Pg:5)

**Supplementary Figure S4.** Impact of free azides/triazole moiety on Rhodamine-B fluorescence. (Pg:6)

**Supplementary Figure S5.** SPAAC reaction for glucosyl azide versus fucosyl azide. (Pg:7)

**Supplementary Figure S6.** *In-vitro* SPAAC reactions to detect inorganic azide, organic azide, and mixed (50% organic+ 50% inorganic) azide mixtures for a fixed total molar basis. (Pg:8)

**Supplementary Figure S7.** SDS-PAGE protein gel for wild-type and mutant TmAfc0306. (Pg:9)

**Supplementary Figure S8.** Chemical rescue activity on pNP-fucose for TmAfc0306 constructs. (Pg:10)

**Supplementary Figure S9.** Thin-layer chromatography results for glycosynthase reaction. (Pg:11)

**Supplementary Figure S10.** *In-vivo* SPAAC reaction and flow cytometry results. (Pg:12)

**Supplementary Figure S11.** Confocal microscopy confirms SPAAC dye permeation into *E. coli*. (Pg:13)

**Supplementary Figure S12.** D224G mutant fucosynthase and wild-type TmAfc fucosidase expressing *E. coli* cell populations characterized by flow cytometry & FACS. (Pg:14)

**Supplementary Figure S13.** Example of fluorescence gates used for FACS sorting of D224G mutant epPCR library. (Pg:15)

**Supplementary Figure S14.** Scatter plots (SSC vs Fls 561nm ex/580/30nm em) for D224G epPCR mutant library supplemented with β-L-fucopyranosyl azide as donor sugar and various distinct acceptor sugars: (A) Galactose, (B) Lactose, and (C) N-acetyl galactosamine. (Pg:16)

**Supplementary Figure S15.** Umbrella sampling histograms for QM/MM simulations. (Pg:17)

**Supplementary Table S1.** Mutagenic primer sequences for performing site directed. (Pg:18)

**Supplementary Table S2:** Primer sequences of vector and insert for error prone PCR. (Pg:19)

**Supplementary Text S1**. Impact of azides and triazoles on Rhodamine-B dye fluorescence. (Pg:20)

**Supplementary Text S2.** Confocal fluorescence microscopy to confirm uptake of DBCO-PEG4-FLUOR 545 into *E. coli* cells. (Pg:22)

**Supplementary Text S3.** Error Prone PCR (epPCR) conditions and validation. (Pg:23)

**Supplementary Text S4.** Protein sequences for wild-type TmAfc0306, D224G mutant, and novel glycosynthase mutants identified with improved activity on pNP xylose. (Pg:25)

**Supplementary Methods.** (Pg:27)

**Supplementary Discussion.** (Pg:32)

**Author Contributions and Conflicts of Interest.** (Pg:35)

**Supplementary References.** (Pg:36)

#### Supplementary Figures

**Supplementary Figure S1. Molecular structure of DBCO-PEG4-Fluor 545**. In Figure 1 the fluorophore is depicted as a colored star symbol. Fluor 545 or tetramethylrhodamine (TAMRA) moiety structure is shown below in red dotted lines.

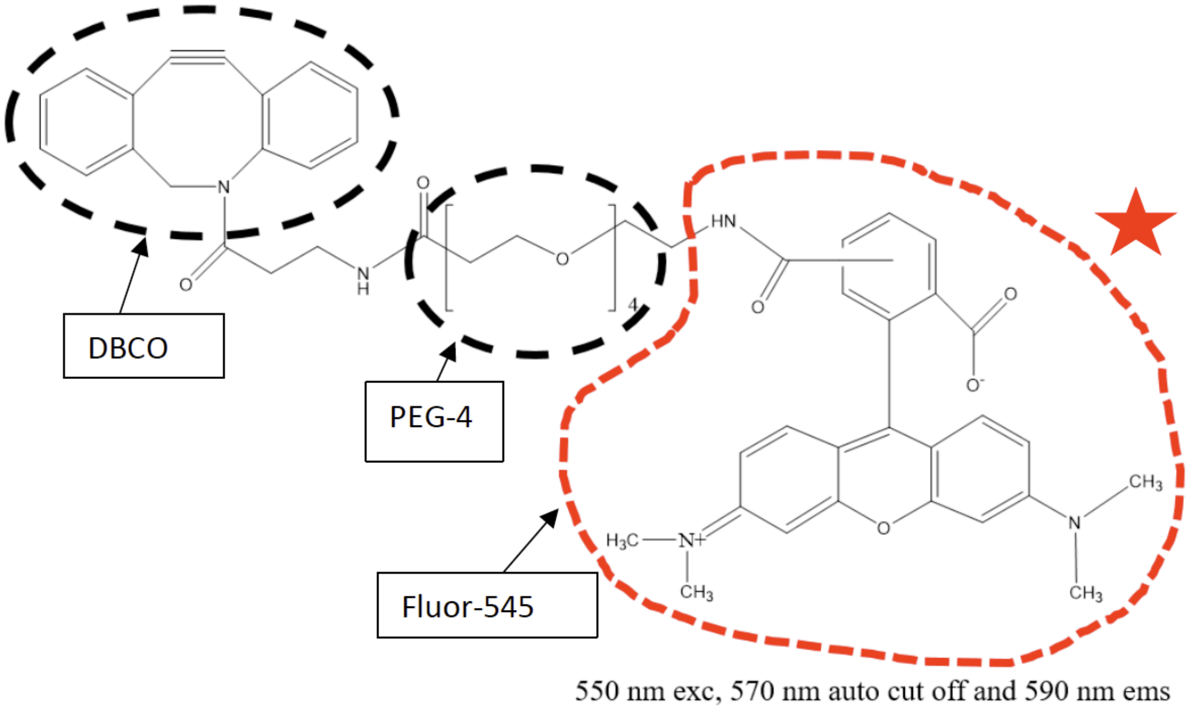

**Supplementary Figure S2. *In-vitro* SPAAC chemistry reaction rates at 25°C (A) versus 10°C (B).** The reduction in fluorescence rates at 25°C were 0.013± 0.001 $s^{-1}$ and 0.0105± 0.002 $s^{-1}$ for sodium azide and glucosyl azide, respectively. At 10°C, the reduction in fluorescence rates were 0.0107± 0.001 $s^{-1}$ and 0.0067± 0.0005 $s^{-1}$ for sodium azide and glucosyl azide, respectively. The reduction in fluorescence rates at 25°C and 10°C were about 3 times lower than click chemistry reaction rate at 37°C (see supplementary figure S5). Hence, 37°C was selected as the optimal temperature for conducting the click chemistry reaction. Also, since 37°C was also optimal condition for *E. coli* cell growth, this temperature was used for all subsequent *in-vivo* click chemistry experiments. The line traces are shown to aid the reader in following the relative change in values. Error bars shown here represent one standard deviation from the reported mean value.

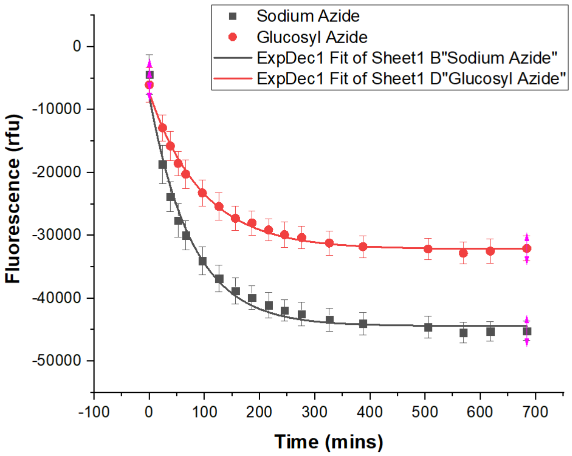


**(A)** Fluorescence decay kinetic curve above is for the reaction between DBCO-PEG4-FLUOR 545 (200 μM) with sodium azide and β-D-glucopyranosyl azide (400 μM) in 1X PBS buffer pH=7.4 at 25°C. Data was collected at 550 nm excitation, 570 nm auto cut off, and 590 nm emission. Table below depicts model fitting analysis results to measure rate of fluorescence decay.

| Model | ExpDec1 | |
|---|---|---|
| Equation | y = A1*exp(-x/t1) + y0 | |
| Plot | Sodium Azide | Glucosyl Azide |
| y0 | -44400.0 ± 384.9 | -32165.4 ± 196.7 |
| A1 | 36419.8 ± 1410.0 | 24817.1 ± 473.3 |
| t1 | 76.9 ± 5.2 | 95.5 ± 3.8 |
| Adj. R-Sq | 0.982 | 0.995 |

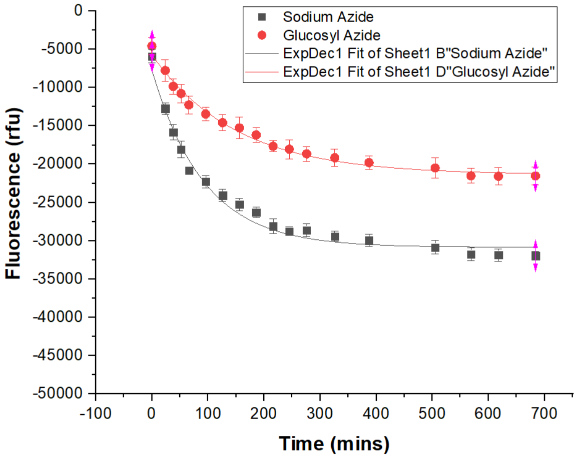


**(B)** Fluorescence decay kinetic curve above is for the reaction between DBCO-PEG4-FLUOR 545 (200 μM) with sodium azide and β-D-glucopyranosyl azide (400 μM) in 1X PBS buffer pH=7.4 at 10°C. Data was collected at 550 nm excitation, 570 nm auto cut off, and 590 nm emission. Table below depicts model fitting analysis results to measure rate of fluorescence decay.

| Model | ExpDec1 | |
|---|---|---|
| Equation | y = A1*exp(-x/t1) + y0 | |
| Plot | Sodium Azide | Glucosyl Azide |
| y0 | -30854.9 ± 453.6 | -21373.8 ± 329.0 |
| A1 | 23172.1 ± 1001.4 | 15670.2 ± 440.1 |
| t1 | 93.1 ± 8.4 | 150.6 ± 10.8 |
| Adj. R-Sq | 0.97 | 0.99 |

**Supplementary Figure S3. Emission fluorescence at varying excitation wavelengths for SPAAC reaction products**. Triazole product formed after DBCO-PEG-Fluor545 reacts with azide gave a maximum difference in 590 nm emitted fluorescence at 550-560 nm excitation wavelength. The line traces are shown to aid the reader in following the relative change in values. Error bars shown here represent one standard deviation from the reported mean value from two biological replicate samples.

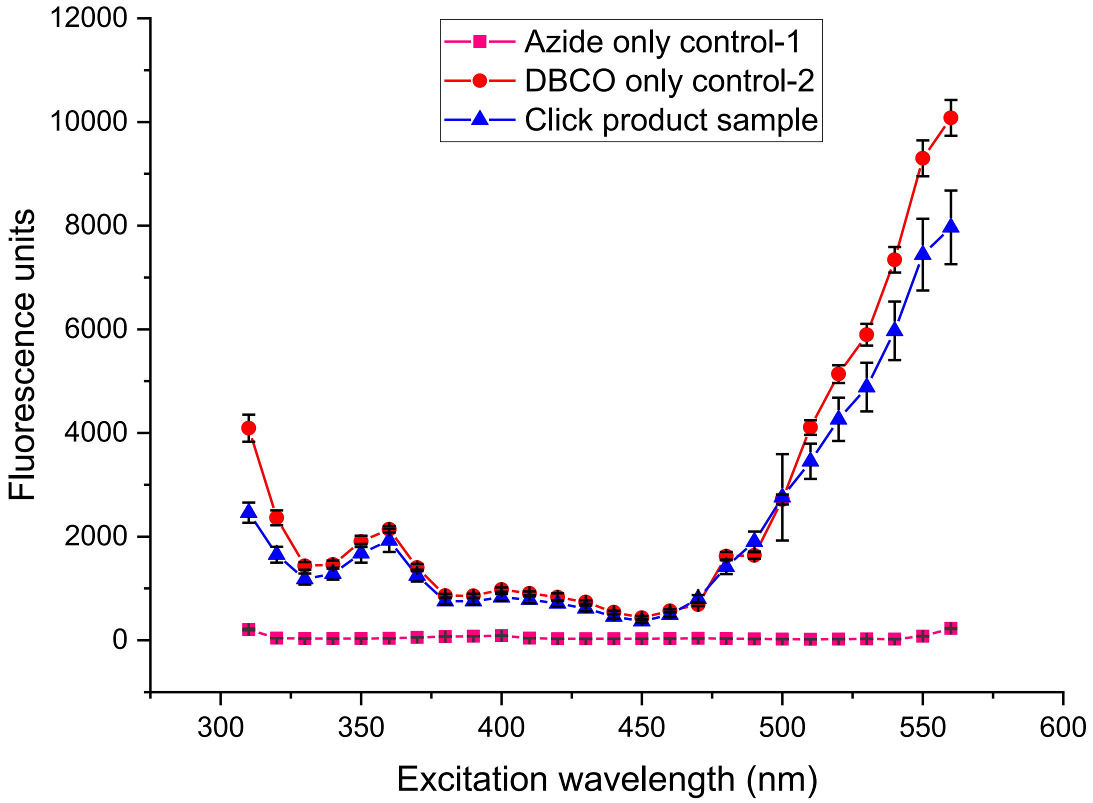

**Supplementary Figure S4. Impact of free azides and triazole moiety on Rhodamine-B fluorescence.** (A) Comparing the molecular structures of Fluor-545 moiety and close structural homolog Rhodamine-B. (B) Relative fluorescence traces for mixtures of Rhodamine-B (200 µM) with sodium azide or β-D-glucopyranosyl azide (400 µM) in 1X PBS buffer pH=7.4 at 37°C measured at 550 nm excitation and 590 nm emission (570 nm auto cut off). Note here that data for Rhodamine-B control alone has been subtracted from all reported data points. (C) UV spectra showing the loss in absorbance at 309 nm upon reaction of DBCO-NHS with sodium azide versus β-D-glucopyranosyl azide at 37°C confirms formation of triazole product using DBCO-NHS moiety. (D) Next, DBCO-NHS SPAAC reaction products first formed in (C) with either organic or inorganic azides that resulted in formation of either glycosylated versus non-glycosylated triazole products were then mixed with Rhodamine-B at time t=0. However, there was no major change in the Rhodamine-B dye fluorescence added exogenously to either preformed triazole products. The line traces are shown to aid the reader in following the relative change in values. Error bars shown here represent one standard deviation from the reported mean value.

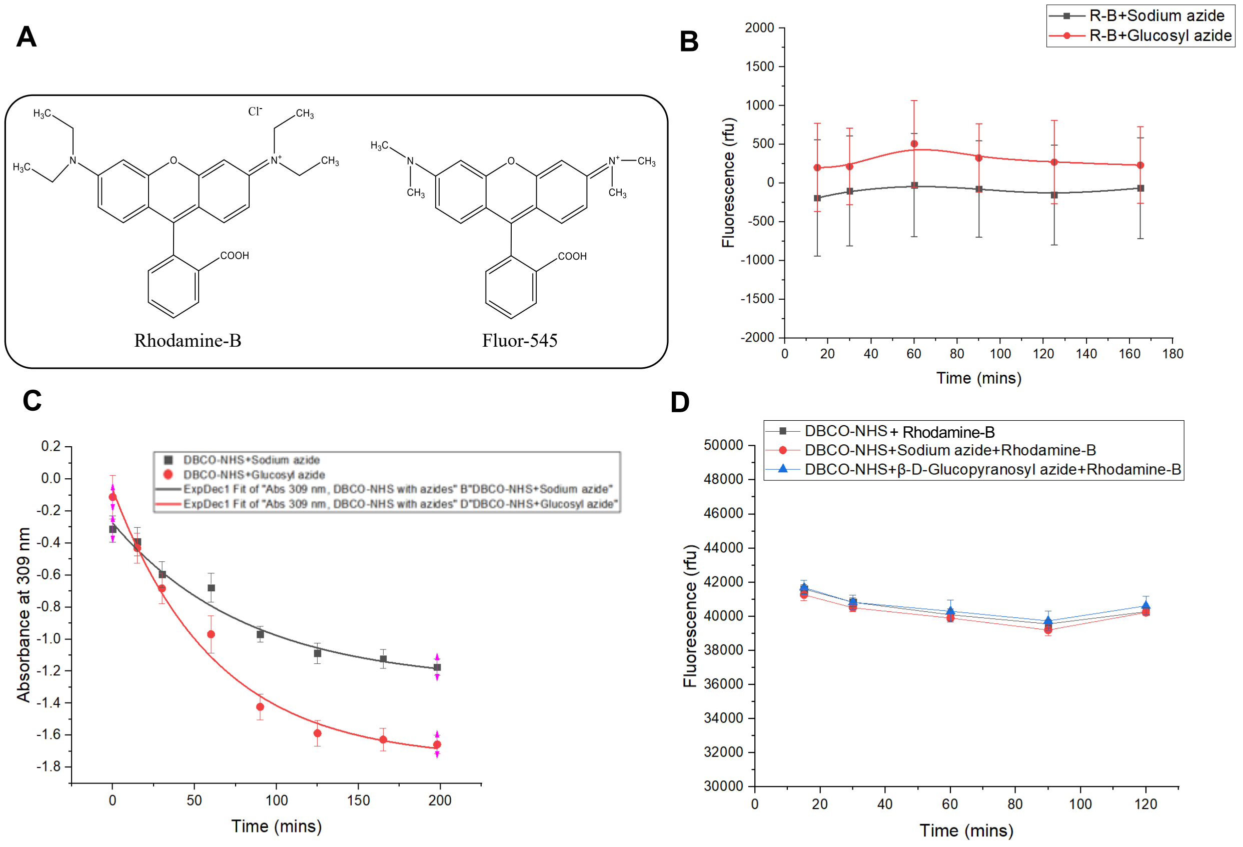

**Supplementary Figure S5. SPAAC reaction for glucosyl azide versus fucosyl azide**. Click chemistry reaction was performed at 37°C using DBCO-PEG4-Fluor 545 with glucosyl azide and fucosyl azide for ~320 minutes total reaction time. We found that click reaction with glucosyl azide versus fucosyl azide showed a very similar drop in fluorescence that was expected due to similarity in the molecular structures of the formed triazole product. Molecular structures of β-D- glucopyranosyl azide and β-L- fucopyranosyl azide shown below. This triazole moiety is expected to be clearly distinct from the one formed from inorganic azides and hence that could explain why the sugar azides behave similarly based on the change in fluorescence associated with the Fluor545 group. The line traces are shown to aid the reader in following the relative change in values. Error bars shown here represent one standard deviation from the reported mean value.

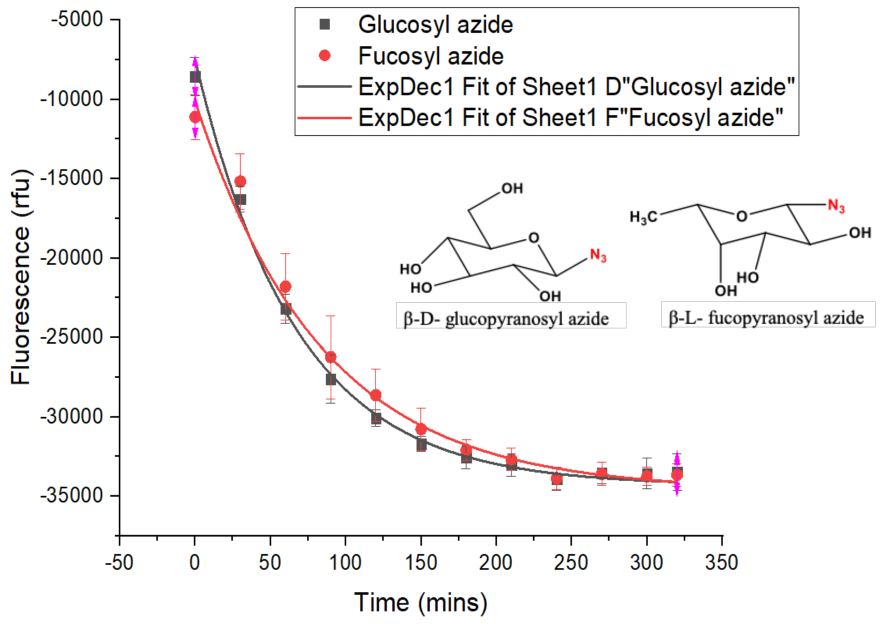


| Model | ExpDec1 | |
|---|---|---|
| Equation | y = A1*exp(-x/t1) + y0 | |
| Plot | Glucosyl azide | Fucosyl azide |
| y0 | -34319.7 ± 290.8 | -34638.0 ± 397.0 |
| A1 | 26808.8 ± 624.6 | 24523.8 ± 851.4 |
| t1 | 67.0 ± 3.4 | 83.9 ± 7.2 |
| Adj. R-Square | 0.99 | 0.99 |

**Supplementary Figure S6. *In-vitro* SPAAC reactions to detect inorganic azide, organic azide, and mixed (50% organic+ 50% inorganic) azide mixtures for a fixed total molar basis.** This experiment was performed to evaluate the effects of click chemistry reaction products on the dye molecule fluorescence at 550 nm excitation, 570 nm auto cutoff and 590 nm emission when an equimolar combination of 50% sodium azide and 50% β-D- glucopyransoyl azide was reacted with DBCO-PEG4-FLUOR 545. It was expected that the fluorescence of click chemistry reaction with a combination of both azides would be lying somewhere between results observed for sodium azide and β-D-glucopyranosyl azide alone. This suggests that if we have mixed concentrations of the organic and inorganic azide formed by glycosynthases of varying degrees of catalytic efficiency, we should still be able to ideally differentiate amongst the products formed and residual substrates left using a similar click chemistry detection assay method. Error bars shown here represent one standard deviation from the reported mean value.

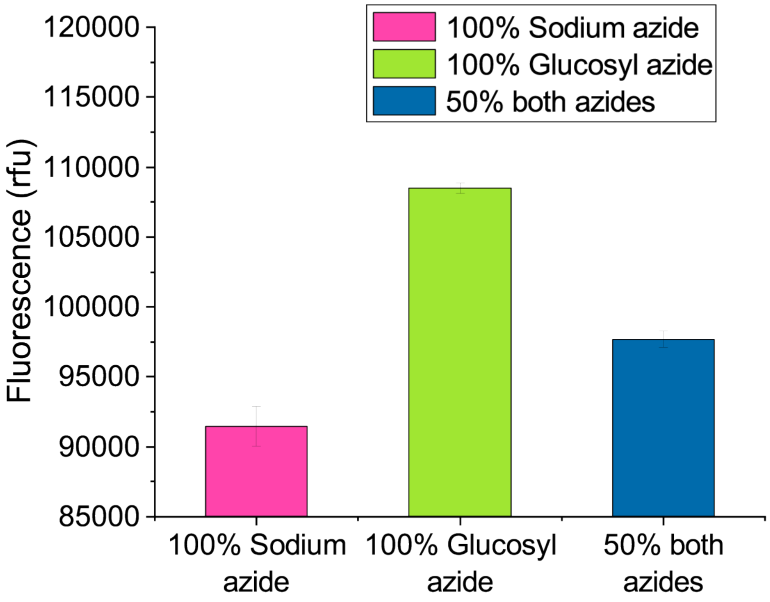

**Supplementary Figure S7. SDS-PAGE protein gel for wild-type TmAfc0306 (WT), mutant D224A TmAfc0306 (A), mutant D224S TmAfc0306 (S), and mutant D224G TmAfc0306 (G) purified proteins used for all reported *in-vitro* glycosynthase reactions**. First lane is the standard protein ladder. Last three lanes correspond to TmAfc0306 mutants (M8, M5, and M9) identified later in this study using the FACS screening methodology and are discussed elsewhere.

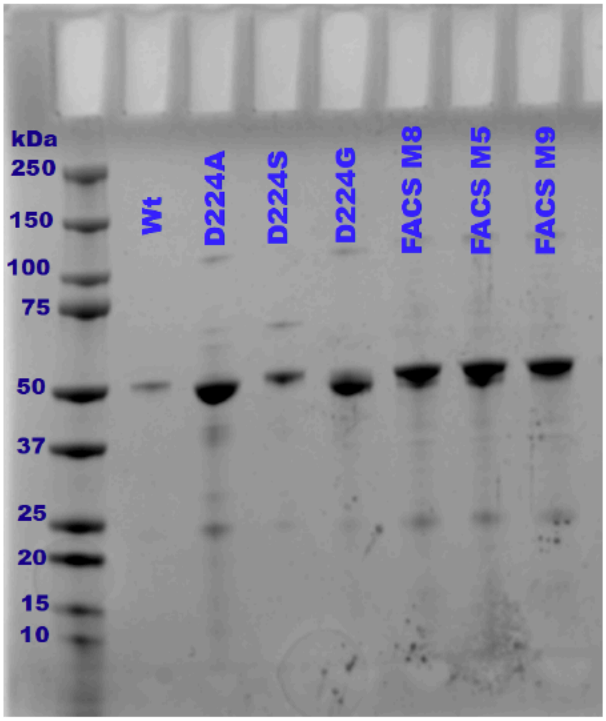

**Supplementary Figure S8. Chemical rescue activity on pNP-fucose substrate for wild-type TmAfc0306 (WT), mutant D224A TmAfc0306 (D224A), mutant D224S TmAfc0306 (D224S), and mutant D224G TmAfc0306 (D224G) purified enzymes in the presence or absence of external nucleophiles is shown here**. Chemical rescue activity assays were carried out in presence and absence of 2M sodium azide and sodium formate as external nucleophiles at 60 °C in 50 mM MES buffer at pH 6.0. Error bars shown here represent one standard deviation from the reported mean value.

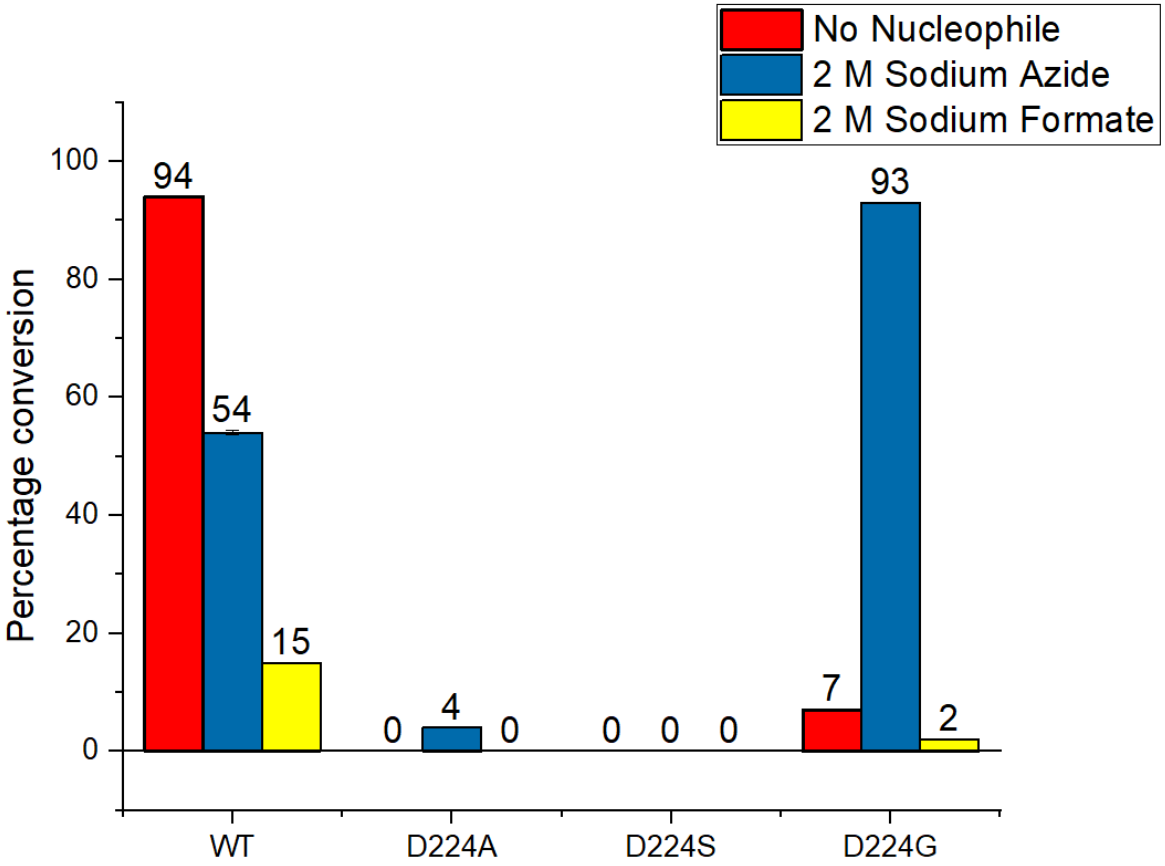

**Supplementary Figure S9. Thin-layer Chromatography (TLC) result for glycosynthase reaction mixtures analysis of substrate/products formed for wild-type (WT) and nucleophile mutant (D224G or Gly) with β-L-fucopyranosyl azide and pNP-β-D-xylopyranoside as donor and acceptor substrates, respectively.** All reactions were conducted in 50 mM MES pH=6.0 at 60 C for 24 hours total reaction time. Here, TLC plate image before start of reaction (left) and after 24 (right) hours reaction time are shown here and all samples were run in duplicates. Glycosynthase reaction products were separated using TLC and visualized using Orcinol reagent. Briefly, the orcinol visualization reagent was sprayed on the TLC plate and then incubated at 100 °C for 15 minutes. The TLC plate was then imaged in a Gel Doc EZ Imager under epi-illumination using white light to perform densitometric analysis of the charred spots observed using the visualization agent. Here, we observe that one of the potential glycosynthase reaction products cannot be clearly observed after Orcinol staining (unlike the UV image taken prior to the Orcinol staining step). This is because residual β-L-fucopyranosyl azide and the formed glycosynthase reaction product/s have similar TLC retention factors. Thus, after Orcinol staining, the glycosynthase reaction product is hidden behind the dominating spot of residual unreacted β-L-fucopyranosyl azide. We also observed minor hydrolysis taking place for the donor sugar (β-L-fucopyranosyl azide) as evident from the presence of fucose.

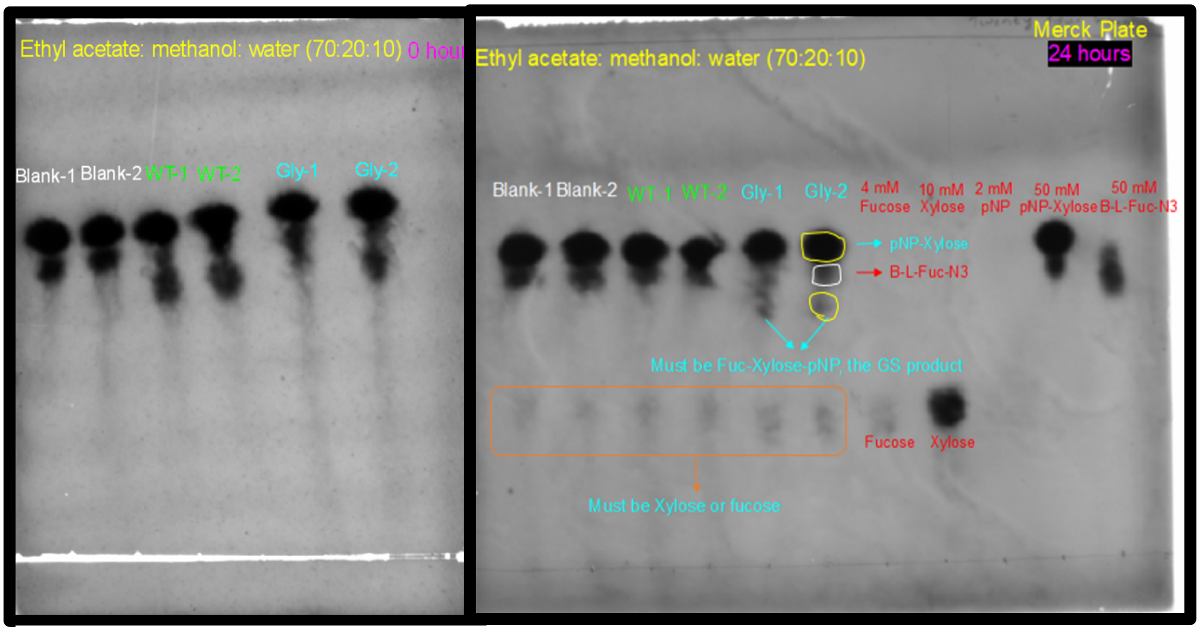

**Supplementary Figure S10. *In-vivo* SPAAC reaction and flow cytometry results summary to provide proof-of-concept data showcasing how cells containing organic and inorganic azide based SPAAC triazole products give differences in relative fluorescence signal (analogous to *in-vitro* results discussed before)**. Click chemistry reaction between DBCO-PEG4-Fluor545 (alkyne; called DBCO in the figure below) and Azide (NaN3 or Glc-N3) was performed *in-vitro* at 1:2 ratio at 37°C for 4 hours. The click chemistry reaction mixture was next incubated with 500 µl of *E. coli* cells at OD=1 for 1 hour. Samples were then run using a flow cytometer (Beckman Coulter CytoFLEX Cytometer) to characterize single-cell fluorescence and the overall cell population distribution. Blue laser was used for excitation (488 nm) and red fluorescence channel filter was set at 585/42 nm BP. Here, data from two independent flow cytometry runs per sample (biological replicates) were used for subsequent analysis. A total of 10,000 events per sample run were captured using flow cytometer and the median *in-vivo* fluorescence observed for all replicate sample runs is reported below. For gating, control cells incubated with sodium azide and glucosyl azide alone were taken as control cells and the fluorescence obtained from the cells is excluded. Error bars shown here represent one standard deviation from the reported mean value for two biological replicate runs.

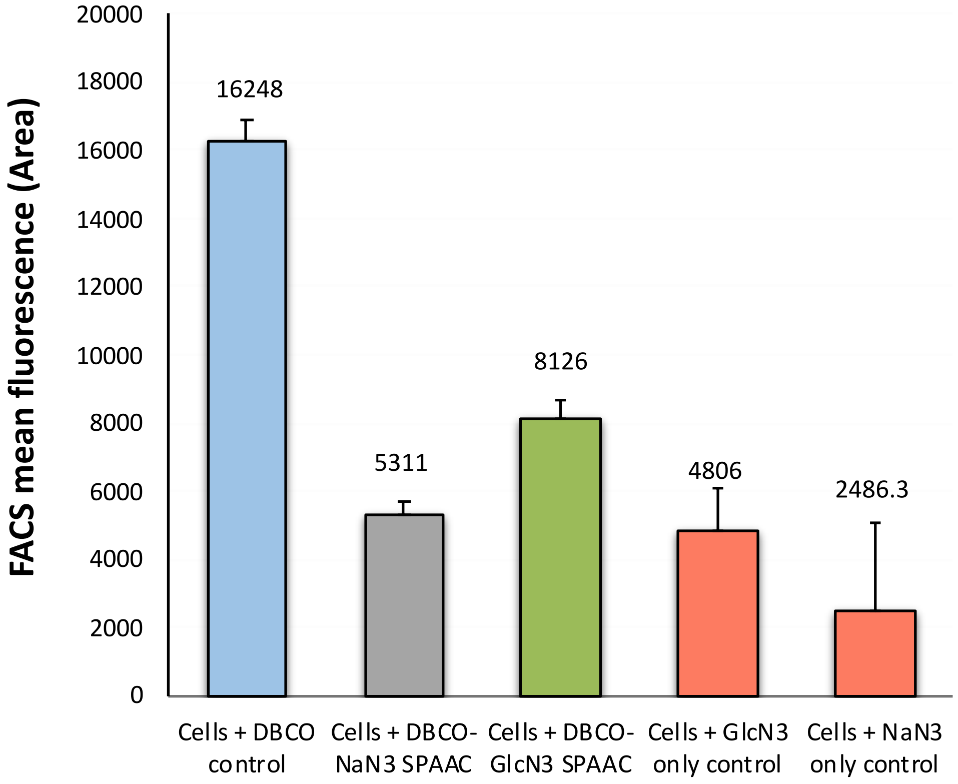

**Supplementary Figure S11. Confocal microscopy confirmed DBCO-PEG4-FLUOR 545 dye can readily permeate inside *E. coli* cells**. (A) See Supplementary Text S2 for details on how these experiments were setup as briefly summarized in figure below. (B) Representative images at various magnifications of unwashed cells confirm the co-localization of the dye (red) with the host DNA (blue) inside the bacterial cells (greyscale).

**A**

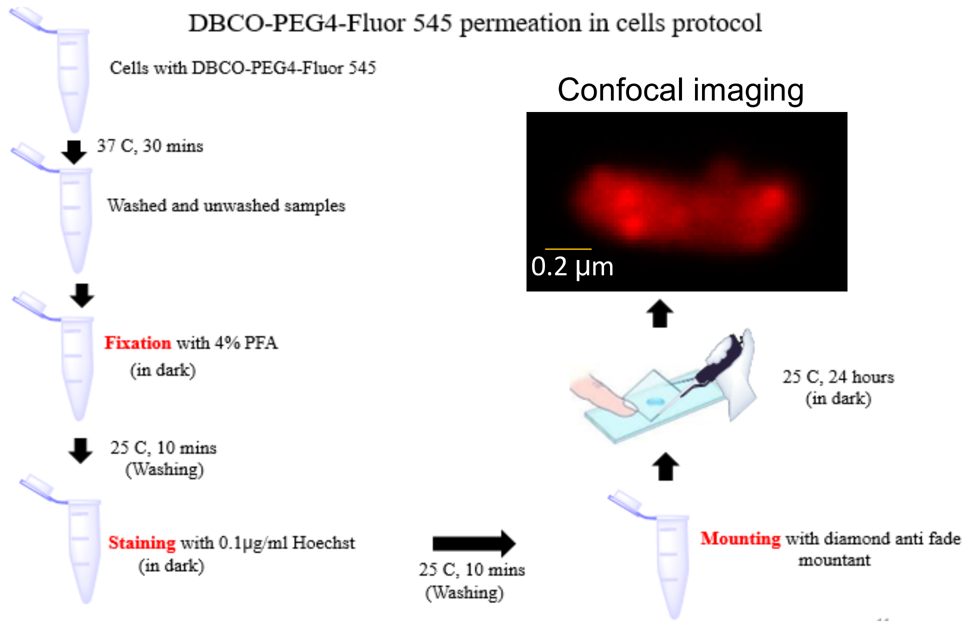


Protocol for sample preparation for confocal microscopy

**B**

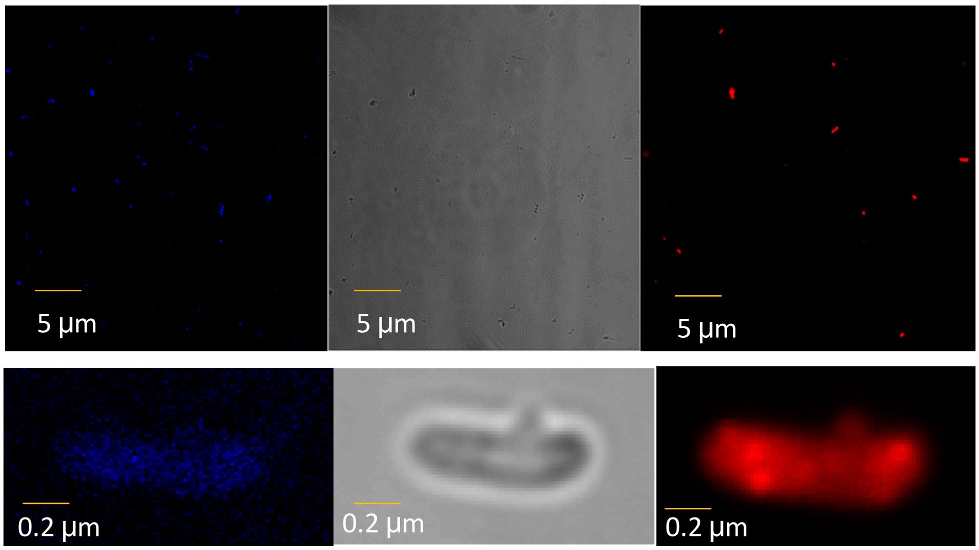


Hoechst staining (0.1 µg/ml) shows DNA co-localization

Bright field Microscopy shows *E. coli* cells at low/high magnification

DBCO (0.1 mM) found co-localized inside *E. coli* confirms uptake

**Supplementary Figure S12. Model fucosynthase (D224G) and wild-type (WT) TmAfc fucosidase enzyme expressing *E. coli* cell populations characterized by flow cytometry (top) & FACS (bottom).** Flow cytometry (Guava EasyCyte) was done using 488 nm excitation and 583 nm emission filters, while FACS (MoFlo Cell Sorter) was done using 488 nm excitation and 575 nm emission filters. This experimental data provided a proof of concept *in-vivo* validation for difference in signals obtained for an active glycosynthase vs. an inactive enzyme control (WT) using both a flow cytometer and FACS instruments. Although the difference in signal was marginal due to the poor activity of D224G (see bottom histogram for total cell count as a function of fluorescence).

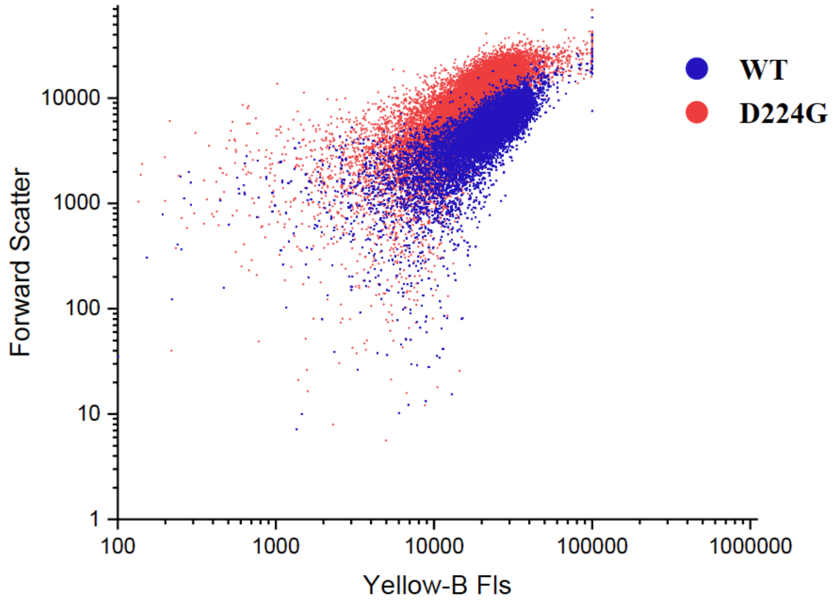


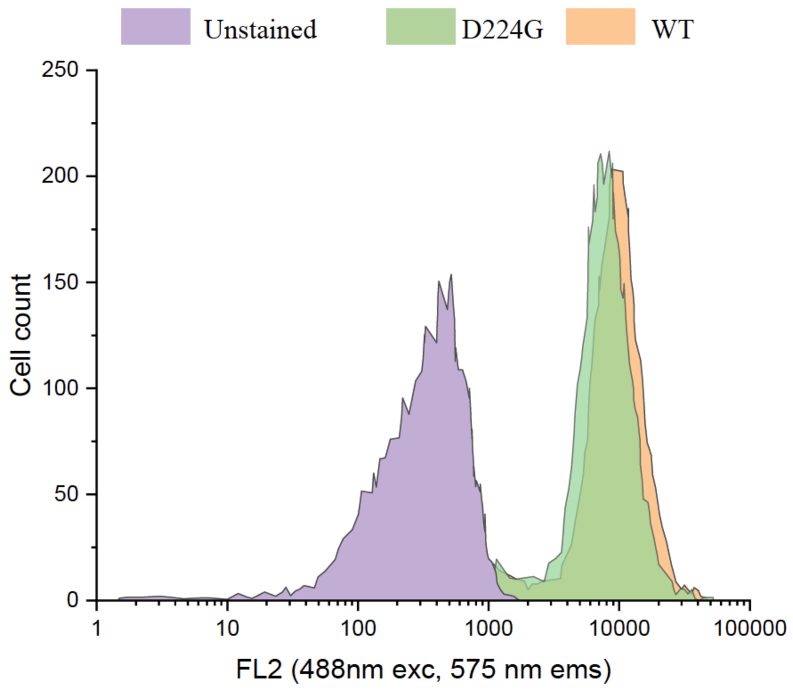

**Supplementary Figure S13. Example of fluorescence gates used for FACS sorting of D224G mutant epPCR library is shown below**. FACS (MoFlo Cell Sorter) was done using 488 nm excitation and 575 nm emission filters. The population of cells which showed low fluorescence signal on x-axis were gated as Gate 1 (or 'Low') and cells with higher fluorescence were gated as Gate 2 (or 'High')**.** Note that the Gate 2 population shows up in the same fluorescence range that overlaps with the fluorescent signals seen for wild type or even D224G cells. However, the clear shift in the fluorescence intensity populations in Gate 1 are indicative of the likely presence of improved glycosynthase mutants from the epPCR library.

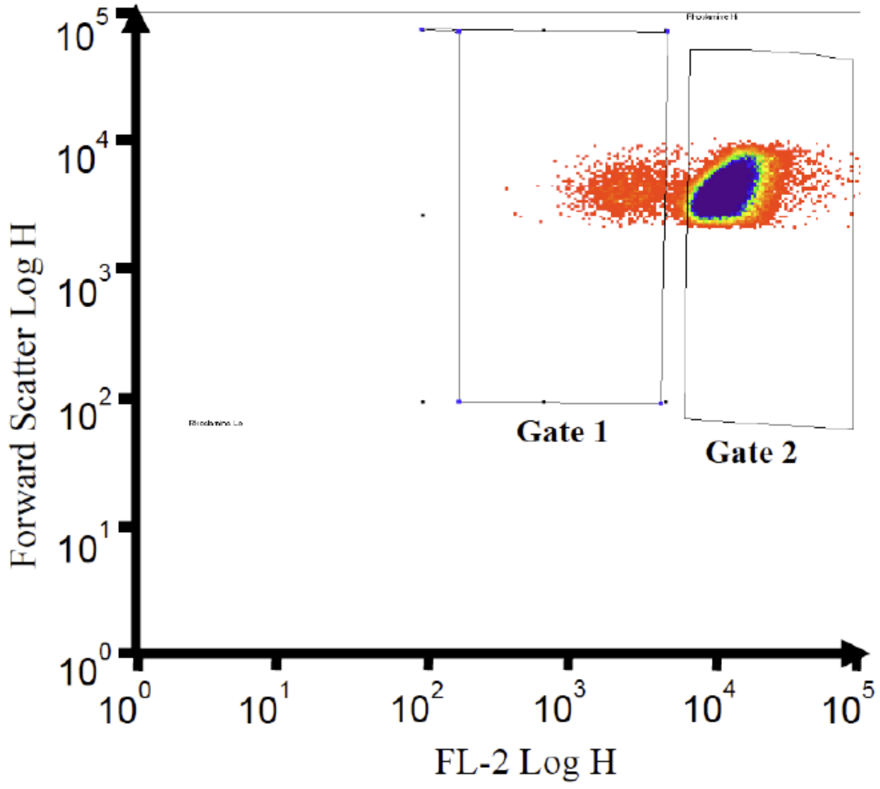

**Supplementary Figure S14. Scatter plots (SSC vs Fls 561nm ex/580/30nm em) for D224G epPCR mutant library supplemented with β-L-fucopyranosyl azide as donor sugar and various distinct acceptor sugars: (A) Galactose, (B) Lactose, and (C) N-acetyl galactosamine.** The gates "Low" and "High" are indicated as rectangles. The cell population percentage in each gate is indicated by the number above the rectangle. Here, we can clearly see a nearly 10-fold significant increase of mutant cells identified in the low gate (~17% of total population) compared to the starting D224G control shown in Figure 4 (1.7% of total population).

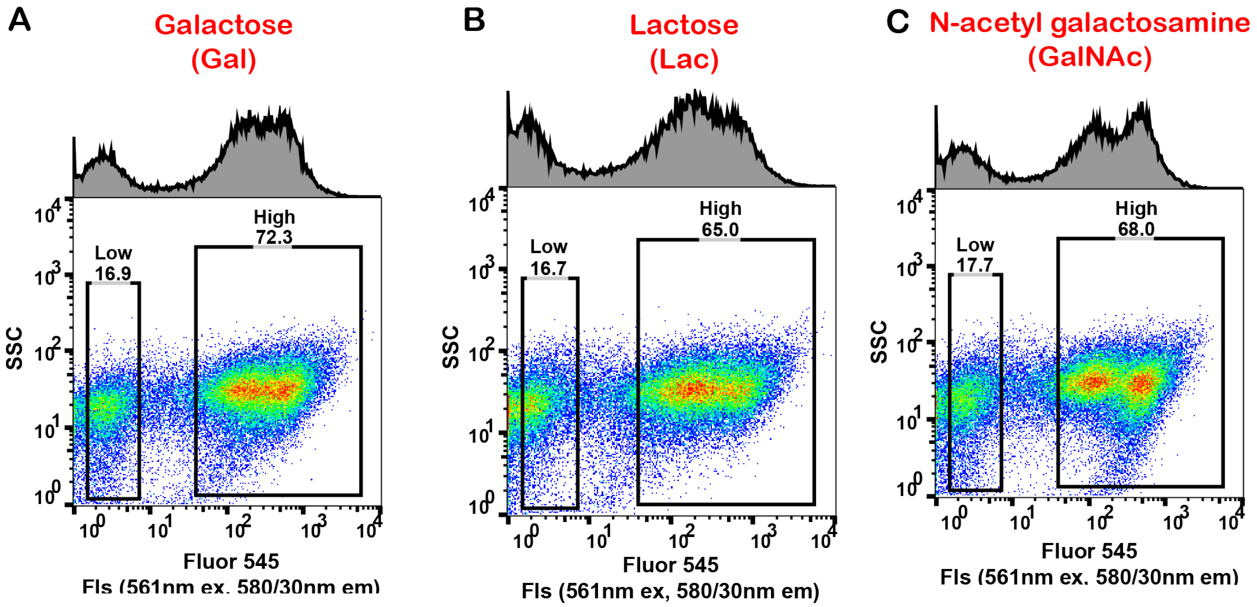

**Supplementary Figure S15. Umbrella sampling histograms for QM/MM simulations.** Histograms showing the sampling along the reaction coordinate from each window in the umbrella sampling procedure described in the "Molecular modeling and simulations". Each histogram represents a different window center, with colors to help distinguish between adjacent windows. The overlap between each pair of adjacent windows ensures sampling along the entire reaction coordinate without any gaps.

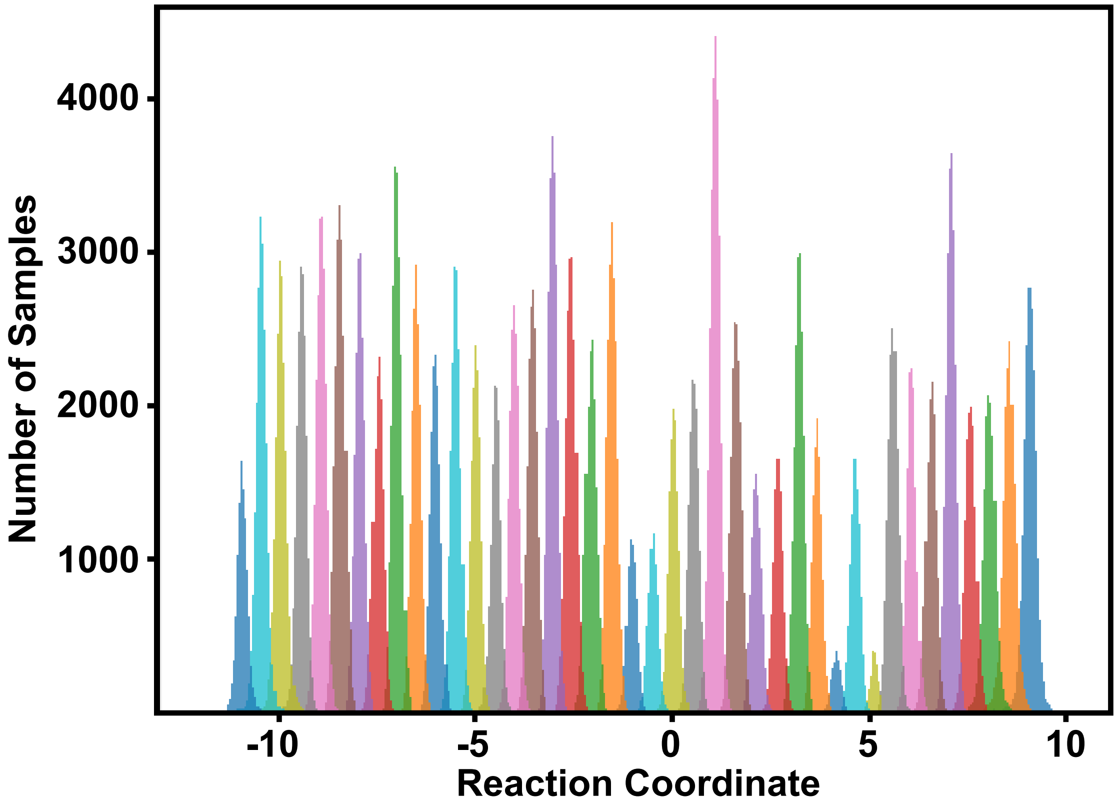

#### Supplementary Tables

**Supplementary Table S1.** Mutagenic primer sequences for performing site directed mutagenesis to generate D224A, D224S, and D224G mutants.

| Primer name | Primer sequence | Melting temperature |
|---|---|---|
| Tm0306_D224A_Forward | GATGTTCTGTGGAACGCCATGGGTTGGCCGGAG | 69 °C |
| Tm0306_D224A_Reverse | CTCCGGCCAACCCATGGCGTTCCACAGAACATC | 69 °C |
| Tm0306_D224S_Forward | GATGTTCTGTGGAACTCCATGGGTTGGCCGGAG | 67.3 °C |
| Tm0306_D224S_Reverse | CTCCGGCCAACCCATGGAGTTCCACAGAACATC | 67.3 °C |
| Tm0306_D224G_Forward | GATGTTCTGTGGAACGGCATGGGTTGGCCGGAG | 69 °C |
| Tm0306_D224G_Reverse | CTCCGGCCAACCCATGCCGTTCCACAGAACATC | 69 °C |

**Supplementary Table S2:** Primer sequences of vector and insert for error prone PCR

| **Primer name** | **Primer sequence** | **Melting temperature** |
|---|---|---|
| Tm0306_WT_epPCR_Vector_Forward | GAATAAGGATCCTCTAGAGTCGAC | 57.5°C |
| Tm0306_WT_epPCR_Vector_Reverse | CATGGCGATCGCCTGG | 57.7°C |
| Tm0306_WT_epPCR_Insert_Forward | CCAGGCGATCGCCATG | 57.7°C |
| Tm0306_WT_epPCR_Insert_Reverse | GTCGACTCTAGAGGATCCTTATTC | 57.5°C |

#### Supplementary Text

**Supplementary Text S1**. **Impact of azides and triazoles on Rhodamine-B dye fluorescence.**

Supplementary Figure S4A shows that Fluor-545 (Tetramethyl rhodamine) and Rhodamine-B (Tetraethyl rhodamine) dyes are structurally similar with a minor difference as the methyl groups in Fluor-545 is replaced by ethyl groups in Rhodamine-B. However, since Fluor-545 dye moiety alone is not readily available from commercial sources. It was either tagged with esters, azides, or other functional groups that might interfere with the SPAAC reaction. So, we instead studied the effect of free azide or SPAAC derived triazole products on the fluorescence of Rhodamine-B dye instead. Our aim here was to study the effect on the fluorescence of the fluorophore moiety alone and the fluorophore moiety in the presence of exogenously added azides or pre-formed triazole products. This experiment was conducted to explore why the Fluor-545 moiety fluorescence differentially reduces upon completion of the SPAAC reaction for organic versus inorganic azides.

*Effect of exogenous free azides on the Rhodamine-B fluorophore group*: There are no previous studies that have tested fluorescence of Fluor 545 (or Rhodamine-B equivalent) alone in the presence of azides. So, this control experiment was conducted to check if free azides alone affect the fluorophore in some way that might cause a reduction in the fluorescence of the final reaction mixture. To test the effect of exogenously added free azide on the fluorophore group, 200 µM Rhodamine-B was mixed with 400 µM of azides (sodium azide and β-D- glucopyransoyl azide) in 1X PBS buffer pH=7.4. Also, 200 µM Rhodamine-B in 1X PBS buffer pH=7.4 without azides was taken as the Rhodamine-B only control. Finally, 400 µM azides in 1X PBS buffer pH=7.4 without Rhodamine-B were taken as the azide only controls. The reaction was incubated at 37°C for 3 hours and the fluorescence spectra for the solution was recorded every 30 minutes at 550 nm excitation, 570 nm auto cutoff and 590 nm emission in a UV spectrophotometer SpectraMax M5e. Respective azides were mixed with Rhodamine-B and the mixture fluorescence was recorded at various time points at 550 nm excitation, 570 nm auto cutoff and 590 nm emission using UV spectrophotometer Spectra Max M5e. From Supplementary Figure S4B we can clearly observe that there is no significant change in Rhodamine-B fluorescence in the presence of either organic or inorganic azides. Therefore, the free azide in the reaction mixture does not likely interact with the fluorophore group to cause a reduction in fluorescence upon formation of the final triazole product.

*Effect of exogenous free triazole products on the Rhodamine-B fluorophore group*: This experiment was performed to check if the triazole moiety formed during the SPAAC reaction impacted the Fluor545 (or Rhodamine-B equivalent) fluorescence. Initially, DBCO-NHS was mixed with respective azides (sodium azide and β-D-glucopyranosyl azide) to form respective SPAAC products. Here, DBCO-NHS had no fluorophore group attached to it unlike the original DBCO-PEG4-Fluor545 reagent. DBCO-NHS controls without azides and azide controls without DBCO-NHS were also taken in the Matrical-plate and incubated at 37°C for 200 minutes. To test the effect of triazole moiety on the fluorophore group, 200 µM of DBCO-NHS was first mixed with 400 μM azides (sodium azide and β-D- glucopyransoyl azide) in 1X PBS buffer pH=7.4 to allow the SPAAC reaction to take place. Here, 200 μM of DBCO-NHS with 1X PBS buffer pH=7.4 without azides was taken as the DBCO-NHS control. Azides were taken with 1X PBS buffer pH=7.4 without DBCO-NHS as azide controls. Only 1X PBS buffer pH=7.4 was taken as the blank for the reaction. The reaction was incubated at 37 °C for 3 hours at 400 rpm. The triazole product formation was first confirmed during the SPAAC reaction by monitoring the change in absorbance at 309 nm for various time points as shown in Supplementary Figure 4C. Note, we can observe that the rate in absorbance decrease is slightly different for DBCO-PEG4-Fluor 545 (Figure 1B**)** and DBCO-NHS (Supplementary Figure S4C). This is likely due to the presence of the PEG-linker in DBCO-PEG4-Fluor 545. Once the triazole product formation was confirmed to be completed after 200 mins total reaction time, Rhodamine-B dye was added to all the wells including the controls and incubated at 37°C while constantly measuring fluorescence at 550 nm excitation, 570 nm auto cutoff and 590 nm emission using a spectrophotometer SpectraMax M5e for various incubation times ranging from 0-120 minutes from the point of addition of the dye. Here, 200 μM of Rhodamine-B was added to all the wells, mixed well, and further incubated at 37 °C for 2 hours.

From Supplementary Figure S4D, we observe that the fluorescence of the solutions containing the triazole based SPAAC reaction products (for both sodium azide and glucosyl azide) was not significantly different when compared to sample containing only DBCO-NHS. Also, there was no significant difference seen between sodium and glucosyl azide as confirmed by student's t-test analysis for p=0.01. It can be inferred from these results that the PEG linker between the triazole product and fluorophore is likely important and facilitates photophysical interactions between the fluorophore moiety and triazole groups that differentially impacts the fluorescence. It is likely that the presence of a PEG linker localizes the fluorophore and the triazole product in close proximity influencing some unique intramolecular interactions that are not seen for intermolecular interactions of the free fluorophore with a free triazole product. For achieving a maximum Förster resonance energy transfer (FRET) signal it is generally observed that the distance between appropriately structured donor and acceptor fluorophores cannot exceed 10 nm.[1] The difference in triazole structures formed for the inorganic azide versus glycosyl-azide could potentially give a distinct quenching of the emitted fluorescence signal from the Rhodamine-B dye if the triazole is within a certain distance and orientation of the fluorophore. However, more work is needed to clearly understand this photophysical phenomenon.

**Supplementary Text S2. Confocal fluorescence microscopy to confirm uptake of DBCO-PEG4-FLUOR 545 into *E. coli* cells**.

Starter culture was inoculated with *E. coli* BL-21 (DE3) glycerol stock for pEC_Tm0306_WT plasmid in 10 ml LB media with 50 µg/ml kanamycin. Here, 5 ml LB media with 50 µg/ml kanamycin alone was taken as a control. The starter culture and the control were incubated at 37 C for 16 hours. Next, 2.25 ml of the starter culture was transferred to 45 ml minimal media with 45 µl kanamycin and 5 ml minimal media with 5 µl kanamycin was taken in a separate tube as control and incubated at 37 C for 16 hours until OD600 of Tm0306_WT reached about 2. The cell culture was centrifuged at 8,000 rpm for 15 minutes and the supernatant was discarded. The culture was washed thrice with equal amount of 1X PBS buffer pH 7.4 and centrifuged at the same conditions as described above. The washed culture was now re-suspended in same amount of 1X PBS buffer pH 7.4. OD600 was measured again and it was found to be 2 again which remains in consistency with the amount of cells in the culture before the washing step. Now, the cell culture was aliquoted into various sterile micro-centrifuge tubes as described below.

First, 2 tubes (labeled as C1 and C3) were prepared as control for the experiment with 200 µl cells and 200 µl DI water. Another 2 tubes (labeled as C5 and S1) were prepared with 200 µl cells and 66 µl of 0.5 mM of DBCO-PEG4-FLUOR 545. All tubes (C1, C3, C5 and S1) were incubated at 37 C for 30 minutes. The samples were centrifuged at 10,000 rpm for 3 minutes and the supernatants were discarded. Now, 50 µl of freshly prepared 4% paraformaldehyde was added to all the samples, mixed well, and incubated at 37 C for 10 minutes. The samples were centrifuged at the same conditions as described above and the supernatants were discarded. The cell pellets obtained were washed twice with 1X PBS buffer pH=7.4 followed by re-suspending in 266 µl of 1X PBS buffer pH=7.4, mixing well, and centrifuging at 10,000 rpm for 3 minutes and finally discarding the supernatant. Next, 50 µl of 0.1 µg/ml Hoechst 33342 was added to S1, mixed well and incubated at 37 C for 10 minutes. S1 was centrifuged at 10,000 rpm for 3 minutes and the supernatant was discarded. S1 was washed twice with 1X PBS buffer pH=7.4 and the supernatants were discarded. C1, C3, C5 and S1 were re-suspended in 100 µl of 1X PBS buffer pH= 7.4 and mixed well. Next, 2 µl of the samples were mixed with 50 µl mounting media (Prolong diamond antifade mounting agent, Catalog number: P36965, Thermo Fisher Scientific) in PCR tubes and centrifuged to remove bubbles. Finally, 10 µl of the samples were placed on a glass slide covered with transparent glass cover slip and incubated at 25 C for 24 hours in dark and visualized under a confocal microscope. See Supplementary Figure S11.

#### Supplementary Text S3. Error Prone PCR (epPCR) conditions

*epPCR reaction conditions*: For insert PCR, 0.5 µM of forward and reverse primers (indicated in Supplementary Table S2) were mixed with 20 ng of plasmid DNA of Tm0306_WT with 0.2 mM of dATP and dGTP, 1 mM of dCTP and dTTP in a 100 µl total reaction volume. The reaction was performed in 1X Taq buffer with 1.25 U of Taq DNA polymerase. 0.1 mM and 0.5 mM $MnCl_2$ was taken in different tubes with (labeled as I1 and I2) and without (labeled as I3 and I4) 1.5 mM and 7 mM $MgCl_2$. The PCR conditions used for insert PCR product amplification are indicated below.

| Process | Temperature | Time |
|---|---|---|
| Initial Denaturation | 95°C | 60 s |
| Denaturation | 95°C | 30 s |
| Annealing | 60°C | 30 s |
| Extension | 68°C | 3 min |
| Final Extension | 68°C | 5 min |
| Hold | 10 °C | ∞ |
| No. of cycles | 20 | |

For Vector PCR products, 0.5 µM of forward and reverse primers (indicated in Supplementary Table S2) were mixed with 20 ng of plasmid DNA of Tm0306_WT in 1X Phusion Master mix in a 50 µl total reaction volume (labeled as V1 and V2). The PCR conditions used for Vector PCR are indicated below.

| Process | Temperature | Time |
|---|---|---|
| Initial Denaturation | 98°C | 30 s |
| Denaturation | 98°C | 10 s |
| Annealing | 60°C | 30 s |
| Extension | 72°C | 3 min |
| Final Extension | 72°C | 5 min |
| Hold | 10 °C | ∞ |
| No. of cycles | 30 | |

*DNA gel for PCR amplification check, PCR product purification*: Once PCR is complete, 2 µl of the PCR product was mixed with 3 µl PCR water and 1 µl of the Purple loading dye and run in SYBR safe DNA gel alongside 5 µl of DNA ladder at 120 V for 40 minutes. With the remaining PCR products, PCR product purification was performed using PCR extraction kit from IBI Scientific.

*DNA gel for purified PCR product, DNA concentration measurement*: Next, 2 µl of the purified PCR product was mixed with 3 µl of PCR water and 1 µl of Purple loading dye alongside 5 µl of DNA ladder and run at 120 V for 40 minutes. The gel was imaged using Gel Doc EZ Imager. The concentration of the purified PCR products was calculated using the gel image.

*Dpn1 digestion, SLIC and transformation*: Reaction mixtures were prepared for Dpn1 digestion. Next, 100 ng of V1 was taken without insert as a control (Reaction-1), 100 ng of V1 was taken with I1 in the Vector: Insert ratios of 1:2.5, 1:5 and 1:10 (Reactions 2,3 and 4 respectively), 100 ng of V1 was taken with I2 in the Vector: Insert ratios of 1:2.5, 1:5 and 1:10 (Reactions 5,6 and 7 respectively), 100 ng of V1 was taken with I4 in the Vector: Insert ratios of 1:2.5, 1:5 and 1:10 (Reactions 8,9 and 10 respectively) in 1X Cut smart buffer in a 10 µl total reaction volume and were digested using 20U of Dpn1 at 37°C for 1 hour. After DPn1 digestion, 1.5U of T4 DNA Polymerase in NEB buffer 2.1 was added to the PCR reaction mixture in a total reaction volume of 20 µl and incubated at 25°C for 5 minutes for SLIC (Sequence Ligation Independent Cloning). The PCR products were incubated on ice immediately after the SLIC run and transformed into E.cloni 10 g cells and incubated at 37°C for 2 hours. The transformation mixture was plated on LB-agar plate with 50 µg/ml kanamycin and incubated at 37°C for 16 hours. Several colonies were observed on the LB agar plates and colony screening was performed to figure out the right colonies.

*Colony Screening:* For colony screening, 30 random colonies were picked from Insert plate (Reaction 3), 30 random colonies were picked from Insert plate (Reaction 9), 5 random colonies were picked from Vector plate (Reaction 1) and transferred to a PCR plate (PCR plate-1) with 5 µl PCR water and incubated at 95°C for 5 minutes. Also, the tip which was used to pick up a particular colony was transferred to LB media with 50 µg/ml kanamycin and incubated at 37°C for 14 - 15 hours. 1 µl of colony from the PCR plate 1 was added to 0.5 µM NcoI forward (TTGCTTTGTGAGCGGATAAC) and 0.5 µM T7 terminator reverse (GCTAGTTATTGCTCAGCGG) primers. The reaction was performed in 1X Master mix in total reaction volume of 40 µl in PCR Plate-2. After colony screening PCR was complete, 2 µl of the PCR reaction mixture was added to 3 µl PCR water and 1 µl of the Purple loading dye alongside 5 µl of the DNA Ladder and loaded onto a DNA gel and run at 120 V for 40 minutes. The DNA gel was imaged using Gel Doc EZ Imager and the positive colonies were identified. The positive colonies were purified using PCR extraction kit and sent for DNA sequencing. The grown colonies were also sent for DNA sequencing after performing mini-prep plasmid extraction for epPCR mutation rate analysis.

**Supplementary Text S4. Protein sequences for wild-type (Wt) TmAfc0306, TmAfc0306 D224G mutant that was used as template for error prone PCR, and novel glycosynthase mutants identified with improved activity on pNP xylose using HTS method.**

**>> TmAfc-Wt**

MISMKPRYKPDWESLREHTVPKWFDKAKFGIFIHWGIYSVPGWATPTGELGKVPMDAWFFQNP
YAEWYENSLRIKESPTWEYHVKTYGENFEYEKFADLFTAEKWDPQEWADLFKKAGAKYVIPTT
KHHDGFCLWGTKYTDFNSVKRGPKRDLVGDLAKAVREAGLRFGVYYSGGLDWRFTTEPIRYPE
DLSYIRPNTYEYADYAYKQVMELVDLYLPDVLWNDMGWPEKGKEDLKYLFAYYYNKHPEGSV
NDRWGVPHWDFKTAEYHVNYPGDLPGYKWEFTRGIGLSFGYNRNEGPEHMLSVEQLVYTLVD
VVSKGGNLLLNVGPKGDGTIPDLQKERLLGLGEWLRKYGDAIYGTSVWERCCAKTEDGTEIRFT
RKCNRIFVIFLGIPTGEKIVIEDLNLSAGTVRHFLTGERLSFKNVGKNLEITVPKKLLETDSITLVLE
AVEE

**>> TmAfc-D224G**

MISMKPRYKPDWESLREHTVPKWFDKAKFGIFIHWGIYSVPGWATPTGELGKVPMDAWFFQNP
YAEWYENSLRIKESPTWEYHVKTYGENFEYEKFADLFTAEKWDPQEWADLFKKAGAKYVIPTT
KHHDGFCLWGTKYTDFNSVKRGPKRDLVGDLAKAVREAGLRFGVYYSGGLDWRFTTEPIRYPE
DLSYIRPNTYEYADYAYKQVMELVDLYLPDVLWNGMGWPEKGKEDLKYLFAYYYNKHPEGSV
NDRWGVPHWDFKTAEYHVNYPGDLPGYKWEFTRGIGLSFGYNRNEGPEHMLSVEQLVYTLVD
VVSKGGNLLLNVGPKGDGTIPDLQKERLLGLGEWLRKYGDAIYGTSVWERCCAKTEDGTEIRFT
RKCNRIFVIFLGIPTGEKIVIEDLNLSAGTVRHFLTGERLSFKNVGKNLEITVPKKLLETDSITLVLE
AVEE

**>> FACS M5 - TmAfc-D224G-N70D-T392S-D400A**

MISMKPRYKPDWESLREHTVPKWFDKAKFGIFIHWGIYSVPGWATPTGELGKVPMDAWFFQNP
YAEWYEDSLRIKESPTWEYHVKTYGENFEYEKFADLFTAEKWDPQEWADLFKKAGAKYVIPTT
KHHDGFCLWGTKYTDFNSVKRGPKRDLVGDLAKAVREAGLRFGVYYSGGLDWRFTTEPIRYPE
DLSYIRPNTYEYADYAYKQVMELVDLYLPDVLWNGMGWPEKGKEDLKYLFAYYYNKHPEGSV
NDRWGVPHWDFKTAEYHVNYPGDLPGYKWEFTRGIGLSFGYNRNEGPEHMLSVEQLVYTLVD
VVSKGGNLLLNVGPKGDGTIPDLQKERLLGLGEWLRKYGDAIYGTSVWERCCAKTEDGTEIRFT
RKCNRIFVIFLGIPSGEKIVIEALNLSAGTVRHFLTGERLSFKNVGKNLEITVPKKLLETDSITLVLE
AVEE

**>> FACS M8 - TmAfc-D224G-N70D-T392S**

MISMKPRYKPDWESLREHTVPKWFDKAKFGIFIHWGIYSVPGWATPTGELGKVPMDAWFFQNP
YAEWYEDSLRIKESPTWEYHVKTYGENFEYEKFADLFTAEKWDPQEWADLFKKAGAKYVIPTT
KHHDGFCLWGTKYTDFNSVKRGPKRDLVGDLAKAVREAGLRFGVYYSGGLDWRFTTEPIRYPE
DLSYIRPNTYEYADYAYKQVMELVDLYLPDVLWNGMGWPEKGKEDLKYLFAYYYNKHPEGSV
NDRWGVPHWDFKTAEYHVNYPGDLPGYKWEFTRGIGLSFGYNRNEGPEHMLSVEQLVYTLVD
VVSKGGNLLLNVGPKGDGTIPDLQKERLLGLGEWLRKYGDAIYGTSVWERCCAKTEDGTEIRFT
RKCNRIFVIFLGIPSGEKIVIEDLNLSAGTVRHFLTGERLSFKNVGKNLEITVPKKLLETDSITLVLE
AVEE

**>> FACS M9 - TmAfc-D224G-N70D-D400A-T413P-T429P**

MISMKPRYKPDWESLREHTVPKWFDKAKFGIFIHWGIYSVPGWATPTGELGKVPMDAWFFQNP
YAEWYEDSLRIKESPTWEYHVKTYGENFEYEKFADLFTAEKWDPQEWADLFKKAGAKYVIPTT
KHHDGFCLWGTKYTDFNSVKRGPKRDLVGDLAKAVREAGLRFGVYYSGGLDWRFTTEPIRYPE
DLSYIRPNTYEYADYAYKQVMELVDLYLPDVLWNGMGWPEKGKEDLKYLFAYYYNKHPEGSV
NDRWGVPHWDFKTAEYHVNYPGDLPGYKWEFTRGIGLSFGYNRNEGPEHMLSVEQLVYTLVD
VVSKGGNLLLNVGPKGDGTIPDLQKERLLGLGEWLRKYGDAIYGTSVWERCCAKTEDGTEIRFT
RKCNRIFVIFLGIPTGEKIVIEALNLSAGTVRHFLPGERLSFKNVGKNLEIPVPKKLLETDSITLVLE
AVEE

**>> FACS M14 - TmAfc-D224G-N70D-T392S-A366V-K395N**

MISMKPRYKPDWESLREHTVPKWFDKAKFGIFIHWGIYSVPGWATPTGELGKVPMDAWFFQNP
YAEWYEDSLRIKESPTWEYHVKTYGENFEYEKFADLFTAEKWDPQEWADLFKKAGAKYVIPTT
KHHDGFCLWGTKYTDFNSVKRGPKRDLVGDLAKAVREAGLRFGVYYSGGLDWRFTTEPIRYPE
DLSYIRPNTYEYADYAYKQVMELVDLYLPDVLWNGMGWPEKGKEDLKYLFAYYYNKHPEGSV
NDRWGVPHWDFKTAEYHVNYPGDLPGYKWEFTRGIGLSFGYNRNEGPEHMLSVEQLVYTLVD
VVSKGGNLLLNVGPKGDGTIPDLQKERLLGLGEWLRKYGDAIYGTSVWERCCVKTEDGTEIRFT
RKCNRIFVIFLGIPSGENIVIEDLNLSAGTVRHFLTGERLSFKNVGKNLEITVPKKLLETDSITLVLE
AVEE

**>> FACS M15 - TmAfc-D224G-N70D-T392S-I428T**

MISMKPRYKPDWESLREHTVPKWFDKAKFGIFIHWGIYSVPGWATPTGELGKVPMDAWFFQNP
YAEWYEDSLRIKESPTWEYHVKTYGENFEYEKFADLFTAEKWDPQEWADLFKKAGAKYVIPTT
KHHDGFCLWGTKYTDFNSVKRGPKRDLVGDLAKAVREAGLRFGVYYSGGLDWRFTTEPIRYPE
DLSYIRPNTYEYADYAYKQVMELVDLYLPDVLWNGMGWPEKGKEDLKYLFAYYYNKHPEGSV
NDRWGVPHWDFKTAEYHVNYPGDLPGYKWEFTRGIGLSFGYNRNEGPEHMLSVEQLVYTLVD
VVSKGGNLLLNVGPKGDGTIPDLQKERLLGLGEWLRKYGDAIYGTSVWERCCAKTEDGTEIRFT
RKCNRIFVIFLGIPSGEKIVIEDLNLSAGTVRHFLTGERLSFKNVGKNLETTVPKKLLETDSITLVLE
AVEE

#### Supplementary Methods

***Gene Synthesis and Cloning:*** A model GH family 29 fucosidase enzyme from a hyperthermophile *Thermotoga maritima* was chosen for this study, also known as Tm-alpha-fucosidase (TmAfc).[2,3] The native (or wild type) gene Tm0306 (Genbank Accession ID NC_000853; see Supplementary Text S4) that encodes TmAfc was codon optimized for *E. coli* expression and custom synthesized with AsiSI and BamH1 restriction sites specific flanking residues in pUC57 by Genscript Biotech Corporation (Piscataway, NJ). The Tm0306 gene was sub-cloned from Genscript's pUC57 vector into our customized pEC vector (with T5 promotor & Kanamycin selection marker[4]) using standard restriction cloning. The catalytic nucleophile of Tm0306 (D224) was mutated into alanine (D224A), serine (D224S), and glycine (D224G) using standard site-directed mutagenesis protocols. Briefly, 0.5 µM of forward and reverse primers for mutagenesis (see Supplementary Table 1) were mixed with 20 ng of plasmid DNA in a 10 µl reaction volume. The reaction was carried out using 1X Master Mix (Phusion DNA polymerase, 200 µM dNTPs, 1X Phusion HF buffer, 1.5 mM $MgCl_2$) with 5% DMSO and the reaction volume was made up to 10 µl by adding nuclease free PCR water. Amplification was confirmed using DNA gel before the PCR amplified reaction mixtures were digested with 10 U of Dpn1 enzyme (New England Biolabs) at 37°C for 1 hour. The Dpn1 digested mixture was transformed into E. Cloni 10 g competent cells (Lucigen, WI) using the Zymo transformation kit and plated onto LB agar plates with appropriate selection marker (Kanamycin). Several random colonies were selected, plasmid DNA was extracted, and verified by DNA sequencing (Genscript, Piscataway, NJ).

***Protein expression and purification:*** Sequence verified wild type (Tm0306_WT) and corresponding nucleophile mutant (Tm0306_D224A/S/G) DNA plasmids were transformed into *E. coli* BL21 (DE3) competent cells and plated onto LB agar plates with 50 µg/ml kanamycin. Individual colonies were picked to inoculate a 50 ml starter culture of LB media supplemented with kanamycin antibiotic (50 µg/ml) and incubated at 37°C for 12-16 hours. Overnight grown cultures were transferred into 1000 ml LB media containing 50 µg/ml kanamycin and grown at 37°C until the culture density reached an $OD_{600}$ of 0.4-0.8. The protein expression was then induced using 0.5 mM Isopropyl β-D-1-thiogalactopyranoside (IPTG) and cultures were incubated at 25°C for 20 hours. The cell pellets were recovered by centrifugation and stored in freezer until needed. The cell pellets were suspended in lysis buffer (20 mM sodium phosphate, 500 mM NaCl and 20% glycerol, pH: 7.4) in a 1:5 ratio of cells to buffer solution (total weight basis), along with protease inhibitor cocktail (1 µM E-64, 0.5 mM benzamidine and 1 mM EDTA) and lysozyme (10 µg/ml) and lysed by sonication on ice. The lysed pellets were then centrifuged and the cell lysate supernatant enriched in the desired soluble protein was recovered. The N-terminal his-tagged proteins of interest were separated from the other undesired *E. coli* proteins using an IMAC (Ni-immobilized metal affinity chromatography) column using the NGC-FPLC system (Bio Rad, Hercules, CA). Briefly, the Ni-IMAC column was equilibrated with the IMAC binding buffer (100 mM MOPS, 10 mM imidazole, 500 mM NaCl, pH: 7.4). Next, the cell lysate supernatant was loaded onto the column and the IMAC binding buffer was run through the column to remove any non-specifically bound proteins from the column. The protein of interest was next eluted with the IMAC elution buffer (100 mM MOPS, 500 mM imidazole, 500 mM NaCl, pH 7.4). The protein was buffer exchanged using desalting columns

(GE Healthcare, Catalog number: 17-0851-01) into 10 mM of 2-morpholin-4-ylethanesulfonic acid or MES at pH 6. The purified protein concentration was estimated using the Spectradrop UV spectrophotometer (SpectraMax M5e) based on 280 nm absorbance. Purity of all enzymes was confirmed by SDS-PAGE (see Supplementary Figure S7) based on gel densitometric analysis using pre-cast stain-free (Bio-Rad) protein electrophoresis gels.

***Fucosidase activity and chemical rescue assays:*** The activity of the purified enzymes Tm0306_WT, Tm0306_D224A/S/G was evaluated using pNP-F (4-nitrophenol α-fucopyranoside) as substrate procured from Carbosynth Limited. Briefly, 1 µg of protein was added to 2 mM pNP-F added in a reaction buffer containing 50 mM MES pH 6 and incubated at 60°C for 1.5 hours. Blank wells with pNP-F alone were taken as buffer/substrate but without the added proteins as controls. Three replicates were taken for each reaction mixture. After 1.5 hours of the reaction, 100 µl of the reaction mixture was transferred to a transparent 96-well microplate along with 100 µl of 1 M NaOH and the absorbance was measured at 410 nm using a UV/Vis spectrophotometer (SpectraMax M5e) to determine total released pNP absorbance upon substrate hydrolysis. pNP calibration curve was built to find the relationship between the measured absorbance and estimated concentration. In order to recover or 'rescue' the hydrolytic activity of the hydrolytically inactive nucleophile mutants, high concentrations of external nucleophiles like sodium azide and sodium formate (2 M each) were additionally added to reaction mixtures and incubated at 60°C for 2 hours. After the reaction was completed, 30 µl of the reaction mixture was transferred to a transparent 96-well microplate and mixed with 70 µl of DI water and 100 µl of 0.1 M NaOH. The absorbance was measured at 410 nm using a UV/Vis spectrophotometer (SpectraMax M5e).

***Glycosynthase in-vitro activity assays:*** For evaluating the glycosynthase activity of Tm0306_WT and Tm0306_D224G, 40 µg of the protein was added to a mixture of 10 mM β-L-fucopyranosyl azide (Catalog number: 66347-26-0, Chemily Glycosciences) and 50 mM pNP-β-D-Xylose (Carbosynth Limited) and incubated at 60°C for 24 hours in 50 mM MES buffer pH 6.0. Two replicates were taken for each reaction mixture. The reaction mixture was then analyzed using Thin Layer Chromatography (TLC) using Silica Gel 60 F254 TLC plates from Merck. The mobile phase used for TLC was ethyl acetate: methanol: water (at 70:20:10 v/v ratios). Standards were also run on the TLC plate to determine the unknown detected spots in reaction sample based on retention factor ($R_f$) value. The plate was epi-illuminated and directly imaged under UV light at wavelength λ=305 nm to visualize pNP and pNP-containing compounds. The plates were then sprayed with visualization solution containing 0.1% orcinol dye in 10% $H_2SO_4$, then dried and heated at 100 °C for 15 min to visualize reducing sugars and acid-labile sugars (see Supplementary Figure S9).

***Click-chemistry in-vitro reactions:*** For copper-free click reaction, 200 µM DBCO-PEG4-FLUOR 545 (Sigma-Aldrich) and 400 µM of respective azides (sodium azide or β-D-glucopyransoyl azide or β-L-fucopyranosyl azide) were mixed in 1X PBS reaction buffer at pH 7.4. Here, 200 µM of DBCO-PEG4-FLUOR 545 with 1X PBS buffer pH 7.4 without azides was taken as the DBCO-PEG4-FLUOR 545 only control. Also, 400 µM azides with 1X PBS buffer without DBCO-PEG4-FLUOR 545 were taken as the azide only controls for the reaction. Three replicates were taken for each reaction mixture. The reaction was incubated for 5 hours total in 384 wells MatriCal plates (GE Healthcare, 28-9324-02) at the following reaction temperatures for optimization: 37 °C, 25 °C, and 10 °C at 400 rpm and fluorescence was recorded every 30 minutes at 550 nm excitation, 570 nm auto cut off and 590 nm emission using a spectrophotometer (SpectraMax M5e). The progress of the click chemistry reaction was also concomitantly monitored by the disappearance of the characteristic UV at 309 nm to follow the formation of the triazole moiety (see Supplementary Figures S2-S5). We also monitored the linear response range of the decrease in fluorescence of the Fluor545 moiety upon completion of the click reaction using inorganic, organic, and mixed inorganic-organic azides. Briefly, 200 μM DBCO-PEG4-FLUOR 545 was reacted with 100% sodium azide (400 μM), 100% β-D-glucopyransoyl azide (400 μM), and with a mixture combination of 50% sodium azide (200 μM) and 50% β-D-glucopyransoyl azide (200 μM) in 1X PBS reaction buffer pH=7.4. Here, 200 μM of DBCO-PEG4-FLUOR 545 with 1X PBS buffer pH=7.4 without azides was taken as the DBCO-PEG4-FLUOR 545 only control. The reaction was incubated at 25°C for 3 hours in 384 wells MatriCal plates and the plate was read for fluorescence at 550 nm excitation, 570 nm auto cutoff and 590 nm emission (see Supplementary Figure S6).

***Error-prone PCR setup via sequence ligation independent cloning (SLIC) approach:*** For insert PCR, 0.5 μM of forward and reverse primers (see Supplementary Table S2) were mixed with 20 ng of plasmid DNA of Tm0306_WT with 0.2 mM of dATP and dGTP, 1 mM of dCTP and dTTP in a 100 μl total reaction volume. The reaction was performed in 1X Taq buffer with 1.25 U of Taq DNA polymerase. 0.1 mM and 0.5 mM $MnCl_2$ was taken in different tubes with (labeled as I1 and I2) and without (labeled as I3 and I4) 1.5 mM and 7 mM $MgCl_2$. For Vector PCR products, 0.5 μM of forward and reverse primers (see Supplementary Table S2) were mixed with 20 ng of plasmid DNA of Tm0306_WT in 1X Phusion Master mix in a 50 μl total reaction volume (labeled as V1 and V2).

***FACS sorting of epPCR library:*** The error-prone PCR library was generated and validated as detailed in the Supplementary Text S3. The epPCR mixture was run on a DNA gel and the bands were extracted using gel extraction. The epPCR products were purified using the PCR clean-up kit from IBI Scientific. Dpn1 digestion was performed at 37°C for 1 hour and SLIC was performed at 25°C for 5 minutes on the extracted products. The SLIC reaction mixture was transformed into E.cloni 10 g cells and incubated at 37°C for 2 hours in SOC media for recovery. After 2 hours, the transformation mixtures were directly transferred to 5 ml LB media as inoculum and grown at 37°C for 16 hours. Next, 1 ml starter cultures were transferred to 20 ml volume cultures in conical flasks with suitable antibiotics and incubated at 37°C for around 2-3 hours until OD600 reached the exponential phase (OD600=0.4-0.8). Then, 1 mM IPTG was added to the cultures and incubated at 37°C for 1 hour to induce protein expression. OD600 was measured after one hour of IPTG induction and 1 ml of the cell cultures were taken out into a sterile micro-centrifuge tubes and centrifuged twice and the supernatants in each round were discarded. Cells were washed twice with 1X PBS buffer pH=7.4 and then re-suspended in 60 μl of 1X PBS pH 7.4 with 10 mM β-L-Fucosyl azide and 25 mM pNP-Xylose added to makeup a total reaction volume of 150 μl. This solution was then incubated at 37°C for 2 hours for the glycosynthase reaction to take place. After 2 hours, the samples were centrifuged and supernatants were discarded. The samples were then re-suspended in PBS buffer and 50 μM DBCO-PEG4-Fluor 545 was added into the total reaction volume of 150 μl and incubated at 37°C for 30 minutes. After 30 minutes, the samples were centrifuged to remove supernatant. Unstained cell samples and D224G (i.e., template DNA) were also taken as controls. The samples were then re-suspended in 1 ml of 1X PBS buffer pH=7.4, filtered using 40 μm filter and run on a FACS instrument (BD Influx High Speed Sorter) with 561 nm excitation laser.

***HPLC analysis of GS reaction products to calculate GS specific activity:*** GS reactions were performed for D224G and the FACS M5 purified proteins to evaluate their specific activities. Briefly, 300 pmoles of each purified protein was reacted with 1 μmole of β-L-fucopyranosyl azide and 25 μmoles of pNP-β-D-Xylose in a 100 μl reaction volume at 60°C. Distinct reaction mixtures were setup for sampling different GS reaction timepoints (i.e., 2 h, 6 h, 10 h, 16 h, 24 h) and three reaction replicates were used for each time point. After each time point, the tubes were rapidly frozen at -20°C to quench the reaction and stored for HPLC-UV analysis. The HPLC analysis was performed on a Shimadzu HPLC system. Briefly, a mobile phase of 90:10 (Acetonitrile:Water) was run through a HILIC column (Shodex Asahipak NH2P-50; 4E 4.6 x 250mm) until a stable baseline is achieved prior to sample injection. Next, 5 μl of reaction mixture was injected onto the column and all pNP-based products (i.e., pNP-xylose, α-L-Fuc-(1,4)-β-D-Xyl-pNP, and α-L-Fuc-(1,3)-β-D-Xyl-pNP) were detected using a DAD detector at 254 nm and 300 nm absorbance wavelengths. The raw data was acquired and analyzed using Shimadzu LabSolutions software. Three distinct peaks were obtained for substrate pNP-Xylose and both GS products for which their respective peak areas were calculated. The area for pNP-Xylose peaks in blank samples was used to normalize and estimate the concentrations of each product in the reaction samples. The initial product formation rate was calculated using the data for 5% conversion of substrate and normalized with the amount of protein added to determine the specific activity of each protein. A two-sided Students t-test was performed for the specific activities of D224G and FACS M5 protein to compared and evaluate their statistical significance.

***Molecular modeling and simulations:*** The molecular model used here was based on a previously published model for the D224G single mutant of the same enzyme.[5] Molecular mechanics (MM) simulations were performed using the Amber 18 software suite.[6] A transition state structure from the previous study was mutated further to match the M5 construct, minimized over 2500 steps, heated from 100 to 300 K over 30,000 2-fs steps, and finally equilibrated over 5 ns with a restraint in place to keep the substrates in the previously identified transition state. The simulations used an Andersen thermostat with a randomization period of 100 steps,[7] a cutoff distance of 8 Å, and the SHAKE algorithm to restrain bonds with hydrogen atoms.[8]

To prepare the system for umbrella sampling, beginning from the equilibrated MM structure the system was further equilibrated over 100 1-fs steps using combined quantum mechanics/molecular mechanics (QM/MM) simulations with the same QM region from the original study, without restraints. Within the QM region the same 8 Å cutoff was used, but SHAKE was not. Because there were no restraints, the system naturally relaxed into one energetic basin (reactants in this case). From there, gentle restraints with initial weight zero and increasing by 0.025 kcal/mol-$Å^2$ each step were used to guide the substrates to the other basin, and this simulation was run until the substrates reached the defined product state. Then, the trajectory was divided into evenly spaced windows along the reaction coordinate every 0.5 units from -11 to 9 (the reaction coordinate is unitless), with the initial coordinates for that window taken from the frame of the trajectory closest to the window center. Using the rxncore model implement in a modified version of Amber,[9] five independent umbrella sampling simulations were performed on these windows, each with step size 0.5 fs and harmonic restraint weight 20 kcal/mol, were run in each window for between 1,811 and 5,437 steps (average 3670.2) each, of which first 1,500 steps were discarded for equilibration. The free energy profile was constructed using pymbar version 3.0.5 (http://www.github.com/choderalab/pymbar).[10,11] The samples were decorrelated using the pymbar.timeseries.subsampleCorrelatedData function[11,12] to ensure only independent samples were considered.

The M5 construct model was also used to perform five unbiased 10-ns MM simulations (of which the first 2.5 ns of each was discarded for equilibration) and compared to the same number and length of simulations for the single (D224G) mutant system. The average by-residue root-mean-square fluctuations (RMSF) were calculated using pytraj[13,14] and subtracted from one another to produce the ΔRMSF data.

#### Supplementary Discussion

Azido sugars have been used extensively as bio-orthogonal reagents in the last two decades to unravel the inner-workings of the glycosylation pathways of cellular systems both at the single-cell and organismal levels.[15,16] However, azido sugars or glycosyl azides have not yet been used extensively as a reagent for synthesis of glycans. Nevertheless, these reagents offer several advantages over other activated sugar donors like glycosyl fluorides or nucleotide sugars for *in-vitro* chemoenzymatic synthesis of glycans. Glycosyl azides can be readily chemically synthesized using one-pot reactions from unprotected sugar monomers, as well as produced enzymatically at high yields, unlike other activated donor sugars like glucosyl fluorides.[3,17,18] Furthermore, GHs are being rapidly discovered through cheaper sequencing of diverse microbial, microbiome, and metagenomic sources.[19,20] These GHs offer a large selection of enzymes that have not yet been exploited for engineering more effective and highly selective GSs for desired donor sugars. Directed evolution of GSs can be used to increase reaction rate and introduce novel donor sugar or acceptor group substrate specificity.[21] However, currently there are very limited options available for universal HTS methods for rational engineering or cell-based directed evolution of GSs.[22–25] Withers first reported a two-plasmid uHTS method where one plasmid contained the GS gene while the other contained a GH-screening enzyme that only releases a fluorophore from the product of the GS reaction but not the original reactants.[22] Similarly, chemical complementation using a yeast three-hybrid system was used to link GS activity to the transcription of a reporter gene, making cell growth dependent on GS reaction product formation.[26] Both these approaches are highly specific to the individual GS family and have limited applicability to screen for novel GS specificity. The first universal HTS method to screen GS libraries (~$10^4$ mutants/day) using glycosyl fluoride as the sugar donor was a pH based assay.[24] Here, hydrofluoric acid, an end-product of the GS reaction using glycosyl fluorides, was detected by a pH sensitive colorimetric indicator. Also, a chemical probe that reacts specifically to the fluoride anion to generate a weak fluorophore was also used recently to screen small GS mutant libraries (~$10^2$ mutants/day).[23] However, to increase the probability of finding rarer GS mutants, fully-automated uHTS techniques capable of handling much larger mutant libraries ($10^6$-$10^{13}$ mutants/day) are necessary. Furthermore, other drawbacks with the current fluoride detection based HTS methods for GS engineering are: (i) low sensitivity limit for detection of reaction products (0.01-10 mM concentration range) which greatly reduces throughput and makes it challenging to fine-tune selection threshold, (ii) inability to distinguish between desired GS activity oligosaccharide products versus side-reaction products due to self-condensation of donor sugars and particularly due to hydrolysis of glycosyl fluorides due to its poor stability in aqueous conditions (e.g., glycosyl fluorides half-life stability ranges between few hours to days for most α- and β-anomers), and (iii) the lack of a sensitive fluorophore than can readily detect unreacted glycosyl fluoride reactant (or released fluoride product), which overall prevents the use of common cell sorting methods necessary to increase screening throughput.[3,27]

Here, we have now developed, validated, and applied a novel SPAAC or click-chemistry enabled uHTS method for screening large GSs libraries. Specifically, other than this report, there are currently no HTS methods available that facilitate ultrahigh-throughput cell-based screening and directed evolution of GSs using azido sugars as glycosyl donors.[27] As proof of concept, we showcased how our screening technique facilitates rapid fluorescence activated cell sorting of a large library of glycosynthase variants (>$10^6$ mutants) expressed in *E. coli* to identify several novel mutants with increased activity for β-fucosyl-azide activated donor sugars towards desired acceptor sugars, demonstrating the broader applicability of this methodology. One of the major advantages of using glycosyl azides as substrates for GS reactions is that the azide/azido moiety can be selectively conjugated to alkyne-based fluorophore groups using a modified Staudinger ligation or copper-free click-chemistry under reaction conditions compatible with the *in vivo* environment.[15,28–30] Similar approaches can be utilized for cell-free based ultrahigh-throughput screening methods for CAZyme engineering as well.[31] Interestingly, the relative difference in the fluorescence intensity of the glycosylated versus non-glycosylated triazole product influences the observed fluorescence for cells enriched in the corresponding SPAAC reaction products. Recent studies have suggested that the alkyne-derived substituent moiety of a “click” triazole can engage in electronic conjugation with the triazolyl core that may have profound influences on the optical properties of these compounds.[32] Photon reabsorption behavior of such compounds can be vexing for the development of energy efficient light-emitting materials, however, this could be advantageous for producing a composite emission color that facilitates selective detection of differentially substitutes triazole products during SPAAC reactions. The influence of the triazole moiety substitution patterns on the electrochemical and photophysical properties of the donor-acceptor groups has been reported in the literature.[33] However, there are no reports, to the best of our knowledge, on the effect of sugar-substituted triazole groups on the variable fluorescence quenching ability due to likely intramolecular photophysical or FRET type interactions with a red-fluorophore group like TAMRA or Rhodamine-B. Future work must resolve the mechanistic basis for selective reduction in fluorescence of non-glycosylated triazole products to facilitate design of more efficient SPAAC reagents to reduce false positives detection using a similar uHTS approach.

Moracci and co-workers had shown how sugar azides can be used for synthesis of glycans using thermophilic GH29 fucosidase enzyme that are able to accommodate slightly larger leaving groups like azides in the mutated D224G active site, instead of smaller leaving groups like fluorides.[3] However, while the D224G was reported to show some fucosynthase activity, we found that the overall activity of this enzyme is actually quite low. Mayes and Bergin recently investigated the first unbiased transition path sampling study of this GH29 fucosynthase enzyme to reveal a single-step mechanism with oxocarbenium-like transition state.[5] Their study suggested one potential explanation for the poor reaction efficiency observed for the D224G fucosynthase was that there were no nearby residues or water molecules to stabilize the departure of the azide group. This could explain why the D224S mutant gave an inactive fucosynthase unlike the D224G due to the relatively bulky serine side chain that likely further restricts azide group accessibility. We have now successfully isolated several novel fucosynthase mutants with improved activity for β-fucosyl-azide towards pNP-xylose. Interestingly, we have identified N70D as a mutation close to the active site that we hypothesize plays an important role in stabilizing substrate binding. However, distal mutations identified outside the active site could not be easily predicted rationally. Some of these distal site mutations are likely relevant to stabilizing the protein fold and/or impact the oligomeric state of the mutant protein that somehow help increase fucosynthase activity. Two major glycosynthase products α-L-Fuc-(1,4)-β-D-Xyl-pNP and α-L-Fuc-(1,3)-β-D-Xyl-pNP were also formed for the improved mutants as well but at much higher turnover rates than previously seen for the D224G variant. Computational transition state and reaction energetics analysis for D224G suggests the formation of both products, but the overall free energy of the reaction being marginally more favorable for the formation of the α-1,3 product.[5] We suspect the additional mutations identified during uHTS screening could have further stabilized the intermediate oxocarbenium-like transition state by aiding in the expulsion of the azide leaving group to increase GS catalytic efficiency for the formation of both α−1,3 and α-1,4 isomers. Interestingly, some of the mutations identified on M5 construct shifted the reaction equilibrium towards the α-1,3 versus α-1,4 isomers suggesting that the uHTS methodology could be useful to also fine-tune glycosidic bond stereochemistry as well. For example, the D224G mutant also gave a ~55:45% (molar basis) of α-L-Fuc-(1,4)-β-D-Xyl-pNP:α-L-Fuc-(1,3)-β-D-Xyl-pNP, as also reported earlier.[3] However, the M5 mutant gave a glycosynthase product distribution of ~48:52% (molar basis) in favor of the α(1,3) isomer. Closer inspect of the *in-silico* mutated M5 structure revealed a native tryptophan residue (W67) adjacent to the N70D mutation that plays an important role in stabilizing the donor sugar and acceptor sugar via hydrogen-bonding and CH-π stacking interactions, respectively. The residues in the active site of the M5 mutant interacting with the docked substrates are shown in green (**Figure 4C**). However, detailed experimental and QM/MM computational analyses are needed to provide a mechanistic basis for the role of mutations both within and outside the active site on improved GS activity. Additional fucosynthase mutants with order/s of magnitude higher glycosynthase activity than the reported M5 mutant is expected as one further optimizes the FACS 'Low' gate for capturing and isolating novel mutants with even lower fluorescence (NOTE: see $<10^0$ fluorescence intensity scatter plot data in red/green contour displayed outside the 'Low' gate for epPCR library as seen in Figure 4A which suggests likely presence of novel GS mutants with even higher fucosynthase activity than M5), as well as screening additional mutant libraries generated using more advanced mutagenesis methods than epPCR.

Finally, this uHTS strategy is currently being exploited to sort mutant D224G epPCR library to identify novel GSs with altered substrate specificity by changing the acceptor sugar from pNP-xylose to either lactose, N-acetylglucosamine, or galactose (Supplementary Figure S14). Here, we can see a nearly 10-fold increase in the relative percentage of mutant cells identified in the low gate (~17% of total population) compared to the starting D224G control shown in Figure 4 (1.7% of total population). These preliminary results clearly highlight that our proposed screening method can be easily applied to readily evolve and screen GS activity for novel acceptor sugars as well. Fucosylated glycans play a critical role in the selective recognition and metabolism of probiotic gut microbes. Human milk oligosaccharides (HMOs) are composed of a tetrasaccharide backbone that is selectively fucosylated to N-acetylglucosamine and/or galactose mirroring the four Lewis blood group antigens. All fucosyltransferases responsible for HMOs fucosylation are yet to be identified, which limits our knowledge on biosynthesis of HMOs. In particular, D224G showed low GS activity with fucosyl azide and lactose as substrates (data not shown). Furthermore, there have been significant advancements recently in the identification and sequencing of entire gene clusters that control the expression of GHs by probiotic gut bacteria (like *Bifidobacterium longum*) dedicated to bacterial growth on HMOs alone.[34] One of the other major roadblocks remains the limited availability of complex glycan reagents that mimic the 100+ naturally occurring HMO to better understand the underlying mechanisms of action on human health. Identification of evolved and novel GS mutants capable of synthesizing 2'-fucosyl lactose, an essential HMO, along with several other Lewis blood group antigen oligosaccharides is currently under investigation in our laboratory.

**Author Contributions:** SPSC conceived the project. AA and CKB designed wet-lab experiments, collected data, and performed all subsequent data analysis. AA, CKB, and SPSC interpreted final results from acquired data. AA drafted the work and substantively revised it with CKB. TB, YW, and HM performed all QM/MM simulations and TB/HM performed all relevant model results analysis. All authors contributed to writing the manuscript and approved the final version.

**Competing Interests:** AA, CKB, and SPSC have filed a US provisional patent application (No. 62/877,021 filed by Rutgers University on July 22nd 2019) on the click-chemistry based cell screening method and novel sequences of GH29 mutants identified using this screening method.

(25) Hayes, M. R.; Bochinsky, K. A.; Seibt, L. S.; Elling, L.; Pietruszka, J. Development of a

Colourimetric Assay for Glycosynthases. *J. Biotechnol.* **2017**, *257*, 162–170. https://doi.org/10.1016/J.JBIOTEC.2017.02.005.
